# A human omentum-specific mesothelial-like stromal population inhibits adipogenesis through IGFBP2 secretion

**DOI:** 10.1101/2023.05.01.538871

**Authors:** Radiana Ferrero, Pernille Yde Rainer, Julie Russeil, Magda Zachara, Joern Pezoldt, Guido van Mierlo, Vincent Gardeux, Wouter Saelens, Daniel Alpern, Lucie Favre, Styliani Mantziari, Tobias Zingg, Nelly Pitteloud, Michel Suter, Maurice Matter, Carles Canto, Bart Deplancke

## Abstract

Adipose tissue plasticity is orchestrated by molecularly and functionally diverse cells within the stromal vascular fraction (SVF). While several mouse and human adipose SVF cellular subpopulations have now been identified, we still lack an understanding of the cellular and functional variability of adipose stem and progenitor cell (ASPC) populations across human fat depots. To address this, we performed single-cell and bulk RNA-seq analyses of >30 Lin–SVF samples across four human adipose depots, revealing two ubiquitous hASPC subpopulations with distinct proliferative and adipogenic properties but also depot- and BMI-dependent proportions. Furthermore, we identified an omental-specific, high *IGFBP2-* expressing stromal population that transitions between mesothelial and mesenchymal cell states and inhibits hASPC adipogenesis through IGFBP2 secretion. Our analyses highlight the molecular and cellular uniqueness of different adipose niches while our discovery of an anti-adipogenic IGFBP2+ omental-specific population provides a new rationale for the biomedically relevant, limited adipogenic capacity of omental hASPCs.

## Introduction

Our understanding of key adipose tissue (AT) phenotypes, such as turnover and expansion dynamics in response to metabolic alterations, is still limited, especially when it comes to human AT. This is further exacerbated by the fact that these phenotypes vary according to the anatomical location of the respective AT. This is illustrated, for example, by the frequent opposition of the overgrown “metabolically healthy” subcutaneous (SC) AT to the “unhealthy” visceral one. However, the terms “visceral” and “subcutaneous” underlie several finer anatomic locations and, with it, potentially more fine-grained characteristics and links to disease^1^. For instance, while SC AT in the thighs has been considered protective against obesity-related insulin resistance, this is not necessarily the case for upper body SC AT accumulation^2^. In part, this has been proposed to be consequent to the intrinsic ability of different depots to increase their size via the generation of new adipocytes (hyperplasia) and/or via (over)growth of their existing adipocytes (hypertrophy)^3^. In this sense, and while increases in fat cell size are generally the main driver of changes in AT mass^4^, femoral subcutaneous fat, which is specialized to provide long-term nutrient storage, has a higher ability to increase fat cell number compared to abdominal subcutaneous fat^5^. At the other side of the spectrum, intraperitoneal visceral fat — such as the omental (OM) depot, for example — generally enlarges through increases in fat cell size rather than number, consistent with its role in storing and releasing nutrients rapidly and its limited space for growth^5^.

Thus, while it is well-accepted that human ATs from distinct anatomical locations expand differently, little is known about what causes these phenotypic divergences. One attractive hypothesis is that these differences could at least be partially driven by variation in the cellular composition of the stromal vascular fraction (SVF) across depots and, more specifically, of adipose stem and progenitor cells (ASPCs). This hypothesis was initially supported by studies showing that SVF cells from human SC AT proliferated and differentiated more potently than those of visceral fat^6^. More recently, comprehensive single-cell transcriptomic (scRNA-seq) atlases of whole human AT, as well as previously published studies, have provided insights into the heterogeneity of human ASPCs (hASPCs)^7–10^. However, these scRNA-seq studies focused on the two most studied ATs: SC and OM. Hence, similarities and/or differences in hASPC composition beyond the SC and OM depots remain elusive.

Studies in mice confirmed that ASPCs are highly heterogeneous across depots, but can be classified into three major overarching ASPC subpopulations^8, 9, 11–18^. These subpopulations, characterized by the expression of specific cell surface markers, exhibit different functional properties^11^. For example, *Dpp4+* (or *Ly6c*+) cells likely represent adipose stem cells (ASCs), a pool of multipotent mesenchymal stem cells that commit to adipogenesis only when exposed to the right mix of factors. In contrast, *Icam1+* (or *Aoc3+*) cells can be classified as pre-adipocytes (PreAs), showing a lower proliferation capacity and a more committed adipogenic state compared with ASCs. Finally, a subset of cells characterized by high expression of *F3* were termed adipogenesis-regulatory cells (Aregs) due to their ability to regulate the differentiation capacity of other ASPCs^7–9, 11–18^. A similar level of phenotypic characterization of hASPC populations is however still lacking, likely reflecting the challenge of having access to and/or gathering enough human biopsy material. Nevertheless, initial efforts to functionally characterize hASPC subpopulations suggested some similarities to the ones identified in mice, with the *DPP4*+ ASPCs being highly proliferative and less adipogenic than the *ICAM1*+ ASPCs^9^. Together, these findings suggest that mouse and human ASPCs might share similar populations. Yet, to date, no systematic, functional characterization of hASPC heterogeneity and behavior has been performed across several human adipose depots.

Here, we provide a comprehensive overview of gene expression profiles of SVF-adherent cells over 30 human donors in four major human depots: SC, perirenal (PR), OM, and mesocolic (MC) AT, combined with scRNA-seq data on ∼34,000 non-immune (CD45–) and non-endothelial (CD31–) SVF cells (SVF/Lin–). We consistently detected two main hASPC subpopulations that are common to all depots. Our analyses also addressed the transcriptional and functional similarities and differences across these depots, as well as a comparison to the most commonly studied mouse ATs. We found that pro-adipogenic/developmental genes are enriched in SC, non-adipogenic/inflammatory ones in OM, mitochondrial/thermogenic ones in PR, and protein folding/trafficking in MC. Furthermore, we established an isolation strategy to isolate, quantify, and characterize different cellular subpopulations in SC, OM, and PR depots with regard to their adipogenic potential and proliferation abilities, validating two surface markers, CD26 and VAP-1, that enable the enrichment of highly proliferative and highly adipogenic cells, respectively, across all depots. Finally, we focused on resolving the mechanism underlying the lower adipogenic potential of OM-isolated SVF-adherent cells, compared to SC and PR ones. We identified a new and OM-specific cell population that inhibits the adipogenic differentiation of hASPCs and is susceptible to undergoing mesothelial-to-mesenchymal transition. We further linked the observed anti-adipogenic effect of this omental population to the secretion of IGFBP2 and activation of the α5β1 integrin receptor in target cells, and hinted at its biomedical relevance by uncovering a significant correlation between inferred *IGFBP2*+ cell abundance and BMI.

## Results

### Human SVF precursor cells exhibit depot-dependent differences in their in vitro adipogenic potential and transcriptome

To characterize the function of SVF-adherent cells, including hASPCs, across distinct human adipose depots, we isolated cell lines from SC (20 donors), PR (8 donors), OM (19 donors), and MC (4 donors) AT (**Supp. Table 1**). As no consensus exists on the surface markers defining hASPCs, and to avoid biasing our strategy towards potential ASPC (sub)populations, we did not implement any enrichment strategy beyond plating SVF cells and culturing SVF-adherent cells. Once confluent, these distinct AT-derived primary cultures were exposed to an adipogenic cocktail for 14 days (**Figure 1A**, see **Methods**). Subsequent staining for lipid droplets revealed that, in line with previous findings^19^, only SVF-adherent cells from ATs located outside the peritoneal cavity (i.e., SC and PR) are able to form mature adipocytes (**Figure 1B-C**). Conversely, cells isolated from intraperitoneal depots (i.e., OM and MC) barely formed any lipid droplets under adipogenic differentiation conditions (**Figure S1A**). Interestingly, while both SC and PR hASPCs differentiated to a higher extent than intraperitoneal cells, PR lines showed the highest adipogenic potential *in vitro*, particularly when cells were differentiated immediately after isolation (**Figure 1B-C**). However, at longer times/passages, the PR-derived cells reduced their level of adipogenicity to that of SC cells (see **Figure 1C** for lowly passaged cells, **Figure S1B** for highly passaged cells). Furthermore, SC and PR lines showed high inter-individual variation in their ability to differentiate, which is observable as an adipogenic potential gradient for SC and PR lines (**Figure 1D**). In contrast, OM and MC lines were systematically resistant to adipogenic differentiation (**Figure 1D**), while also being the slowest growing lines (**Figure S1C**).

**Figure 1.**
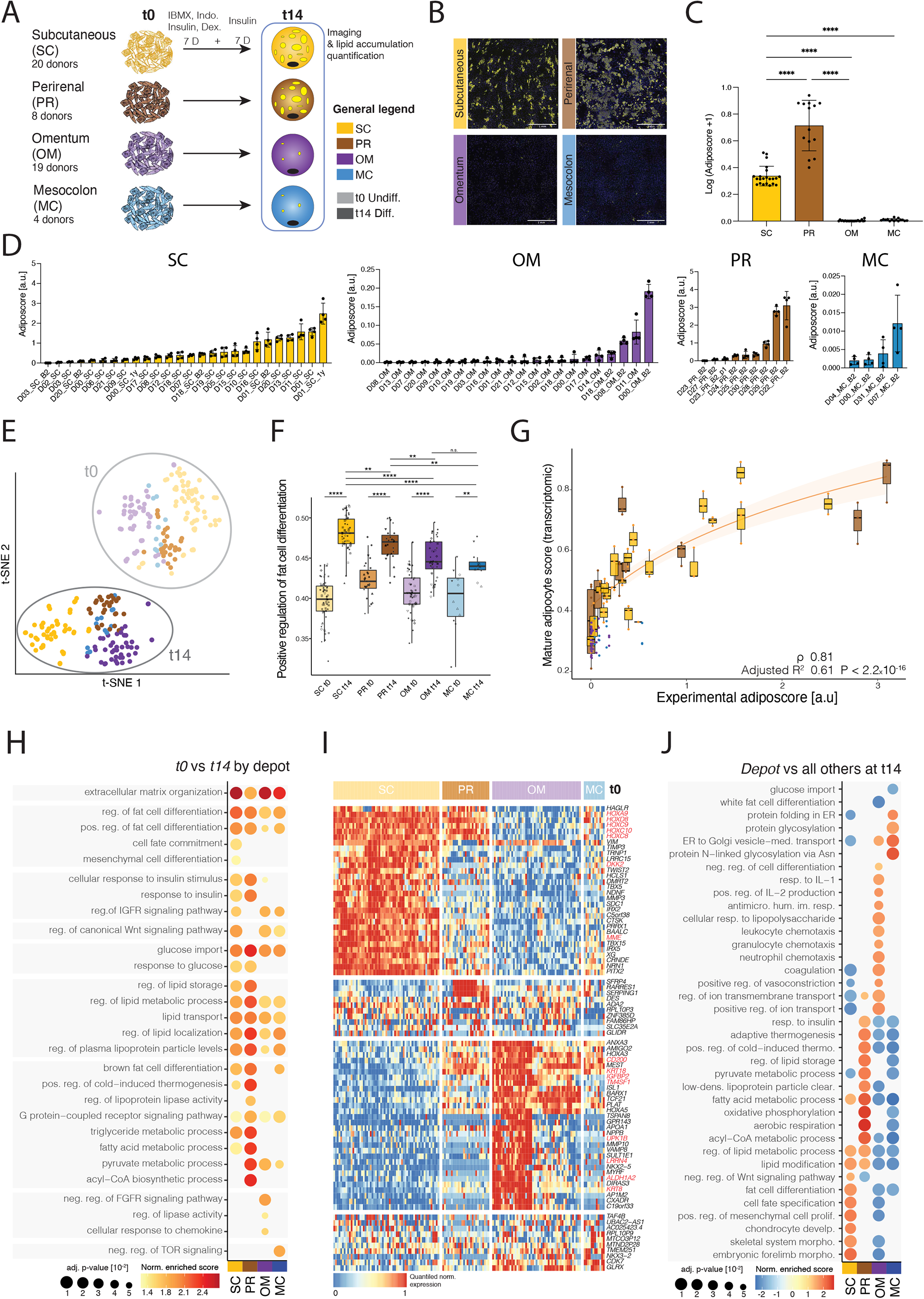
Human SVF cells exhibit depot-dependent differences in their in vitro adipogenic potential and transcriptome **(A)** Scheme of the experimental setup. Primary SVF-adherent cell lines from human subcutaneous (SC), perirenal (PR), omental (OM), and mesocolic (MC) adipose tissues were cultured in parallel and harvested before (t0) or after 14 days of differentiation (t14) for transcriptomic (BRB-seq) analysis; the same lines were seeded in separate assay plates to quantify their adipogenic potential using the adiposcore (see **Methods**). **(B)** Representative fluorescence microscopy images of SVF-adherent cells directly after isolation, expansion to confluence and adipogenic induction (t14); Yellow – Bodipy staining for lipids, blue-Hoechst staining for DNA, scale bar = 1 mm. **(C)** Barplot showing the log(adiposcore + 1) quantification of SVF-adherent cells in **B**; n = 14-22, 4-5 donors, 3-5 independent wells. **(D)** Distribution of the adiposcores (see **Methods**) across all sequenced cell lines from the four depots ordered by increasing values for cell lines with 2 to 6 passages; SC, yellow: n=104, 4 independent wells, 26 cell lines, 20 donors (D); PR, brown: n=36, 4 independent wells, 9 cell lines, 8 donors; OM, purple: n=88, 4 independent wells, 22 cell lines, 18 donors; MC, blue: n=16, 4 independent wells, 4 cell lines, 4 donors; B2 indicates biological replicate from the same individual, and 1y indicates same donor but 1y post-surgery timepoint cell sampling. **(E)** t-SNE map based on the transcriptomic (BRB-seq) data of SVF-adherent cells from the indicated adipose depots (SC – yellow, PR – brown, OM – purple, MC – blue) and time points (t0 light, t14 – dark); n = 12-61, 4-20 biological replicates, 1-4 independent replicates for each. **(F)** Boxplot displaying the “Positive regulation of fat cell differentiation score”, based on the scaled expression of the corresponding GO term (GO:0045600) of the data shown in **E**. **(G)** Scatter plot showing the relationship between the image quantification-based experimental adiposcore in **G** *versus* the “mature adipocyte score” based on the scaled expression of well-known adipogenic markers (see **Methods**) of the transcriptomic samples from the same donor. Samples are grouped by depots and donors. Spearman correlation and adjusted R^2^ of y∼log(x+1) (plotted orange line with 95% confidence interval) values are indicated. **(H)** Dot plot showing enriched, representative terms found by GSEA performed on the differential gene expression analysis results of t0 *versus.* t14 samples for each depot of the data in **E**. **(I)** Heatmap of top differentially expressed genes when comparing the indicated depot *versus* the three others at t0 of the data shown in **E**. (J) Dot plot showing representative, enriched terms found by GSEA performed on the differential gene analysis results of each indicated depot *versus* the others at t14 of the data shown in **E**. \**p* ≤ 0.05, \*\**p* ≤ 0.01, \*\*\**p* ≤ 0.001, \*\*\*\**p* ≤ 0.0001, One-Way ANOVA and Tukey HSD *post hoc* test (**C**), unpaired two-sided *t*-test (**F**).

We explored possible correlations between our experimental adiposcore (**Figure 1D**, see **Methods**) and physiological parameters such as BMI, age, and gender of the donors but found no correlations except for a tendency for PR cells to be less adipogenic in women and elderly people (**Figure S1D**-**H**). However, we acknowledge that our cohort’s demographic characteristics can bias these observations (**Figure S1D**, **Supp. Tables 1** and **2**), as patients were mainly young and obese, and only a relatively small proportion of PR samples could be analyzed (n=8).

To explore if the striking adipogenic difference between intra-peritoneal and extra-peritoneal cell lines is reflected in their respective transcriptomes, we performed bulk RNA barcoding and sequencing (BRB-seq)^20^ of SVF-adherent cells from different individuals and depots, both at the undifferentiated state (t0) and after 14 days of adipogenic differentiation (t14) (SC n=22, OM n=16, PR n=8, MC n=4, **Figure 1A**). We found that the major source of variation is explained by the exposure to the adipogenic cocktail, followed by the anatomic origin of the cell lines (**Figures 1E** and **S1I**-**M**). We observed that all samples at t0 highly express *THY1*, a well-known mesenchymal marker^21^, at similar levels, except OM samples in which it is slightly but significantly lower expressed (**Figure S1N**). The exposure to a differentiation cocktail induced genes related to extracellular remodeling, insulin response, and positive regulation of fat cell differentiation in cells from all depots (**Figure 1F** and **S1O**-**P**). However, most of these adipogenesis-related terms were more enriched in SC and PR compared to OM and MC (**Figure 1F** and **S1O**-**P**). In addition, golden standard markers of adipogenesis and mature adipocytes such as *FABP4*, *PPARG*, *CEBPA*, *ADIPOQ*, *PLIN1-2-4*, *LPL,* and others (see **Methods**) were solely upregulated in PR and SC samples post-differentiation (**Figure S1Q**). The expression of mature adipocyte markers correlated with the lipid droplet accumulation of the corresponding lines as quantified by the image-based adiposcore (ρ=0.81, **Figure 1G**, see **Methods**), showing that inter-individual variability in terms of adipogenicity is also reflected at the transcriptomic level.

Pathway analyses of our transcriptomic data illustrated how programs related to lipid storage and fatty acid metabolism were exclusively enriched in PR and SC-derived cells upon differentiation (**Figure 1H**). Transcriptomic comparisons of undifferentiated cells at t0 revealed that developmental genes such as *HOXC8-10*, *HOXA9*, *and HOXD8* were highly expressed in SC samples (**Figure 1I**), as previously reported^22, 23^. This was further illustrated by the enrichment of numerous terms linked to morphogenesis and development compared to the other depots both at t0 and t14 (**Figures 1I-J**). Interestingly, at t14, SC samples also showed enrichment of (fat) cell differentiation-related terms compared to the other depots, even considering the highly adipogenic PR samples (**Figure 1J**). In contrast, PR-enriched genes at t14 were related to thermogenesis and oxidative metabolism, suggesting that these cells have brown-like or beige-like adipocyte characteristics (**Figure 1J**)^24, 25^. In OM samples, we observed a non-adipogenic gene expression signature with positive and negative enrichment of the terms “negative regulation of differentiation” and “white fat cell differentiation” respectively, compared to cells from the other adipose depots at t14 (**Figure 1J**). Undifferentiated OM cells also exhibited significantly higher expression of genes linked to an inflammatory response, which remained after exposure to an adipogenic cocktail (**Figures 1J** and **S1R**). This is not entirely unexpected given that the OM samples that were analyzed using BRB-seq mainly originated from obese patients undergoing bariatric surgery (**Figure S1D** and **Supp. Table 1**), whose OM fat has previously been reported to show signs of inflammation^1, 26–28^. Interestingly, in both t0 and t14 time points, OM cells showed an enrichment of genes linked to the vasculature and epithelium/endothelium development (**Figures 1J** and **S1S**), suggesting the presence of cells of epithelial nature, and not only mesenchymal ones, in OM SVF-adherent cells. Finally, genes that were specifically expressed in MC compared to other depots were linked to ER stress, protein folding and trafficking (**Figure 1J**).

Taken together, we found that cultured SVF cells from each depot feature specific gene signatures, highlighting the regional specialization of AT based on its anatomical location (**Figure 1J**). In addition, the observed experimental adipogenic potential was mirrored by the up- or down-regulation of pro-adipogenic markers in extraperitoneal and intraperitoneal adipose depot-derived cells, respectively. Finally, mesenchymal markers were highly expressed in SVF-adherent cells from all depots, validating the high enrichment of hASPCs in the SV-adherent fraction (**Figure S1N**). However, OM-derived samples also expressed an enigmatic epithelial gene signature (**Figure S1S** and see below).

### Human adipose-derived stromal cells are highly heterogeneous at the single-cell level

Next, we explored whether the observed transcriptomic and phenotypic differences across depots could in fact be driven by underlying cellular heterogeneity. To do so, we performed scRNA-seq of SVF Lin– (i.e., CD45–/CD31–) cells that were isolated from SC (n=3), OM (n=3), MC (n=2, from the same donor), and PR (n=3) adipose samples (**Supp. Table 3**), analyzing a total of 34’126 cells (on average, ∼8’500 cells per depot). We first analyzed each resulting dataset independently, i.e., per depot and per donor, uncovering high heterogeneity in and between each dataset, as driven by four major subpopulations: two hASPC ones (see below), vascular smooth muscle progenitor cells (VSMPs), and mesothelial cells (**Figure 2A**). We then performed three independent analyses to explore if the identified subpopulations share molecular features across depots and donors. First, we calculated the overlap of the top cluster markers between datasets (**Figure S2A**). We found that, while the percentage of shared markers tends to be the highest within samples isolated from the same depot or donor (**Figure S2B**-**C**), the overlap across depots and donors is, on average, over 50% for most of the identified subpopulations (**Figure S2A**). This result was confirmed when projecting each dataset onto each other using scmap^29^, revealing that on average more than 75% of cells from one specific population projected onto the corresponding population in other datasets, regardless of the depot of origin (**Figure S2D**). Finally, we integrated the data by considering each dataset as a different batch and correcting accordingly. Once again, we observed an excellent overlap of the depot-counterpart populations in the t-SNE space (**Figure 2B**), which was further confirmed by clustering analysis (**Figure 2C**). Focusing on hASPCs, our results indicate that human adipose SVF from four depots, SC, PR, OM, and MC, contains at least two main hASPC subpopulations (**Figure 2D**), characterized by high expression of *THY1* and *PDGFRA* (**Figure 2E**). To explore the universality of this finding, we assessed yet another unexplored AT, namely the AT surrounding the gallbladder in a subset of morbidly obese patients. Even if relatively few hASPCs were ultimately captured, we still retrieved the two main hASPCs subpopulations (n=1, **Figure S2E**).

**Figure 2.**
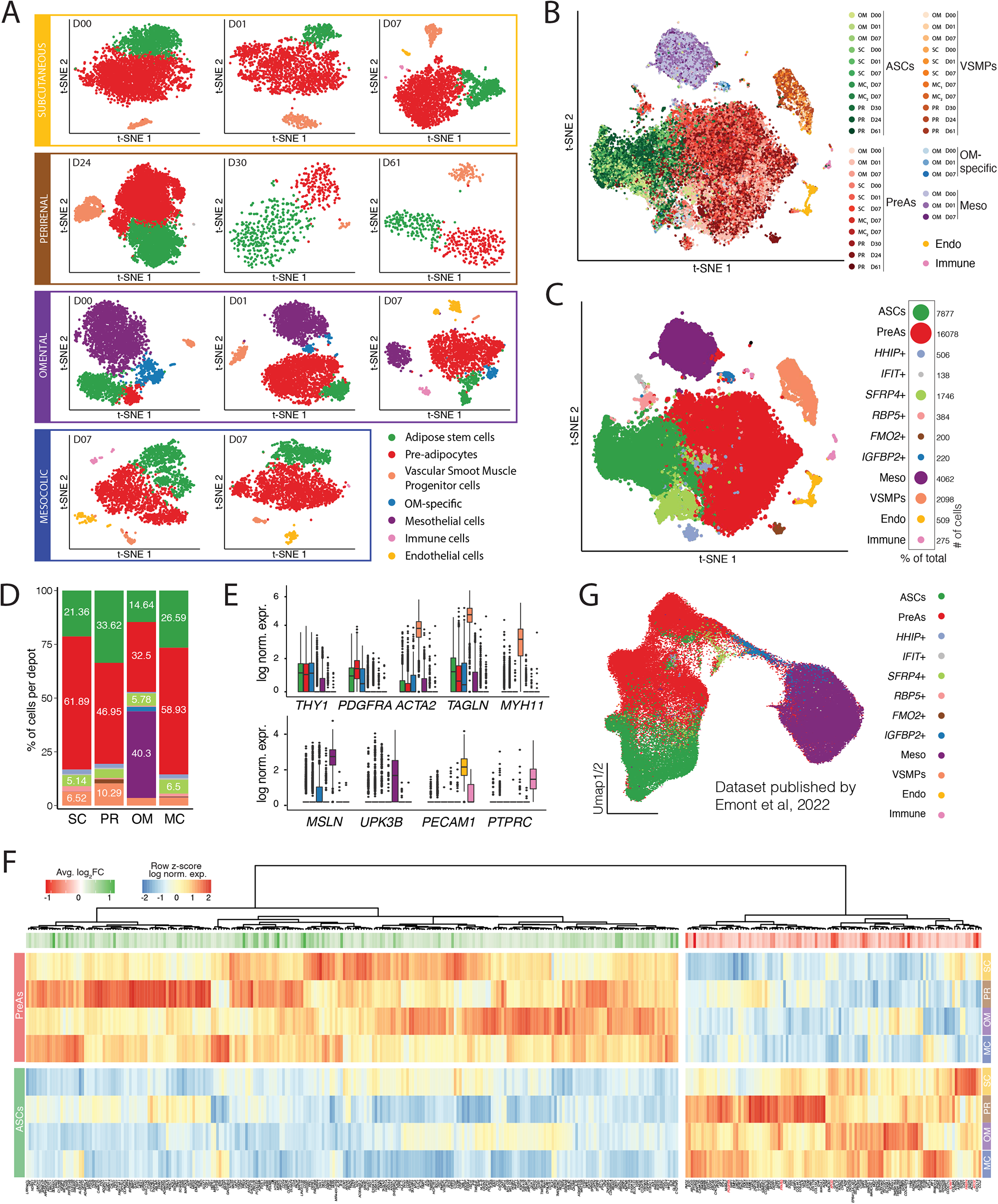
Human adipose-derived stromal cells are highly heterogeneous at the single-cell level. **(A)** t-SNE cell maps of individual scRNA-seq datasets of SVF Lin– cells isolated from four adipose depots (Subcutaneous (SC), omentum (OM), mesocolic (MC), and perineral (PR)) and six different donors (*D*, as indicated in the corner of each t-SNE, see **Supp. Table 3**), visualizing the identified subpopulations of hASPCs following the legend below. The number of cells per dataset from left to right were: SC – 3929, 4169, 2162; PR – 4262, 2042, 2670; OM – 8583, 600, 509; MC – 2650, 2550. **(B)** t-SNE cell map of integrated scRNA-seq datasets across four depots and 6 donors (*D*) (**Supp. table 3**): OM, n=3, SC, n=3, and MC, n=2 (same donor) from matched donors, and PR, n=3, from unmatched donors colored by the clustering of each dataset analyzed individually shown in **A**. **(C)** t-SNE cell map of the data introduced in **B** colored by the identified clustering: Adipose Stem Cells (ASCs) – green, Pre-adipocytes (PreAs) – red, *HHIP*+ ASPCs – light blue, *IFIT*+ ASPCs – gray, *SFRP4*+ ASPCs – light green, *RBP5*+ ASPCs – light-red, *FMO2*+ ASPCs – brown, mesothelial cells (Meso) – purple, vascular smooth muscle progenitor cells (VSMPs) – orange, endothelial cells (Endo) – yellow, and immune cells (Immune) – pink. The percentage of cells belonging to each cluster is shown by a dot plot, with the exact number of cells on the right. **(D)** Bar plot displaying the percentage of cells in each cluster, shown in **C** (excluding immune and endothelial cells) in each depot. **(E)** Box plot showing the log normalized gene expression distribution of selected markers in the different subpopulations depicted in panel **C**. **(F)** Heatmap of the differentially expressed genes between the Adipose Stem Cell (ASC) and the Pre-adipocyte (PreA) populations across depots, based on the data in **C**. **(G)** UMAP of hASPCs and human mesothelial cells from scRNA-seq data published in Emont et al.^8^ colored by the predicted cell type/state when transferring our cell cluster annotation.

Based on their respective gene expression signatures, we labeled those two hASPC subpopulations as adipose stem cells (ASCs) and pre-adipocytes (PreAs) (**Figure 2C**, **F**). Indeed, ASCs from all depots shared a gene signature enriched for *DPP4*, *CD55*, and *PI16*, and showed enrichment in genes involved in proliferation, collagen synthesis and stemness (**Figures 2F** and **S2F**). On the other hand, PreAs differentially expressed known markers of committed adipogenic cells such as *PPARG*, *FABP4*, *PDGFRA*, *APOC*, and *APOE,* and showed enrichment of terms linked to differentiation, commitment, and lipid transport (**Figures 2F** and **S2F**). Furthermore, our annotations are consistent with the two ASPC states observed in human SC AT and predicted for OM AT using independent reference human atlases^8–10^ (**Figures 2G** and **S2G**). To our knowledge, these hASPC states have never been described for human anatomical locations beyond SC and OM.

In sum, we found that, at the single cell level, two canonical hASPC populations – the adipose stem cells and the pre-adipocytes – dominate the transcriptomic landscape of SVF and are retrieved in each analyzed depot. Besides these two, we further detected VSMPs and mesothelial cells together with a number of relatively small clusters that we detail in the next section.

### Common and unique stromal populations exist across adipose depots and their molecular signatures highly overlap with murine counterparts

Next to the ASCs and PreAs, five depot-ubiquitous (VSMPs, *HHIP*+, *IFIT*+, *SFRP4*+, *RBP5*+), one PR and MC-specific (*FMO2*+) and two OM-specific (Mesothelial and *IGFBP2*+) clusters were identified (**Figure 2C**, **D**). All of them were characterized by a unique gene expression signature (**Figure 3A**), which was not always intuitively linked to the adipogenic lineage (e.g. for VSMPs and mesothelial cells).

**Figure 3.**
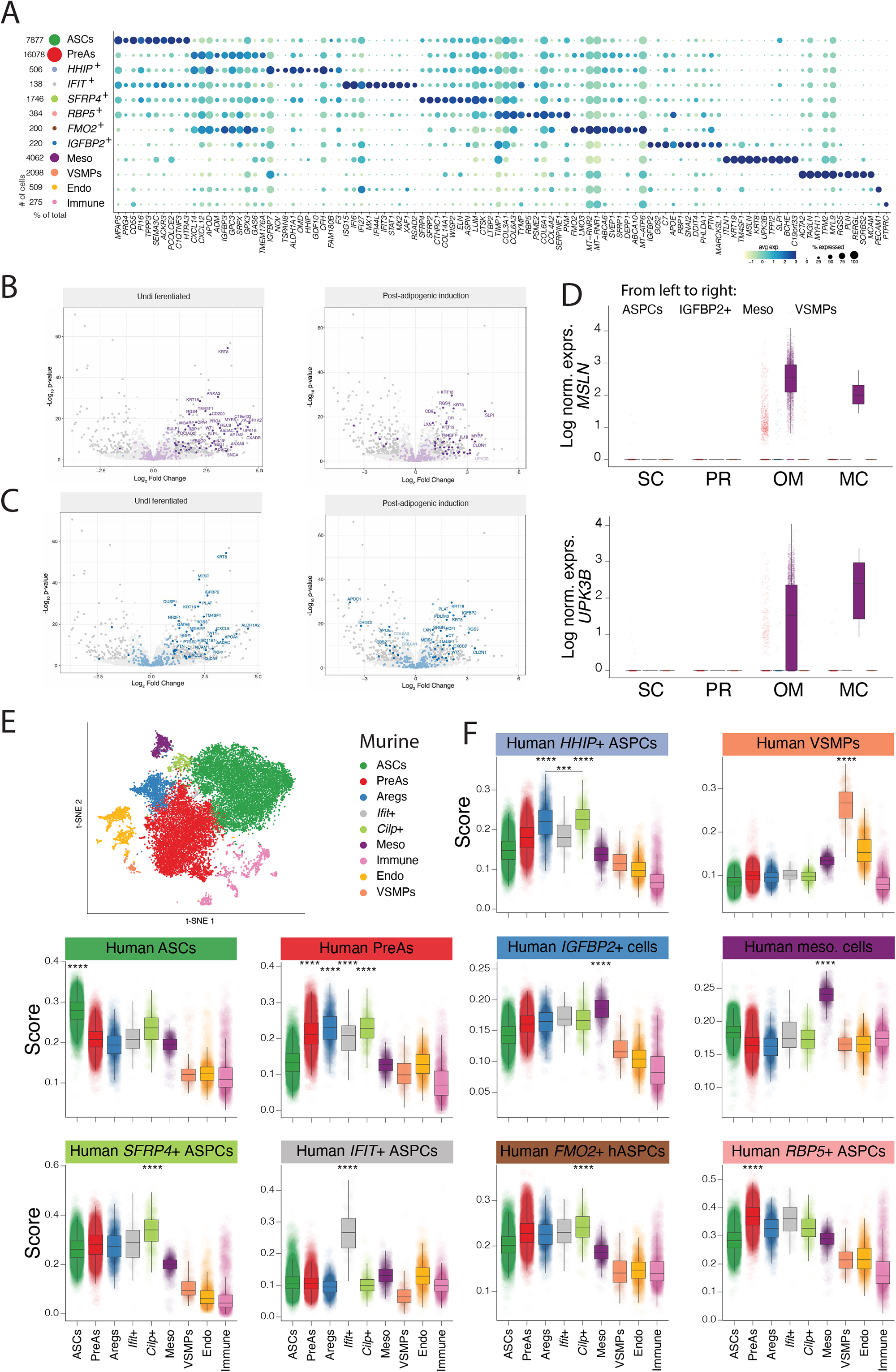
Common and unique stromal populations exist across adipose depots and their molecular signatures highly overlap with murine counterparts. **(A)** Dot plot of 10 markers among the main specific foreach cluster identified in Figure 2C. **(B)** Volcano plot displaying differential gene expression results based on the BRB-seq^20^ data of SVF-adherent cells from the omentum (OM) adipose depot versus SVF-adherent cells from other depots (subcutaneous (SC), perirenal (PR), and mesocolic (MC). The top mesothelial markers identified using scRNA-seq datasets are highlighted in purple, while significantly differentially expressed genes (log2FC > 1, adjusted p-value < 0.01) are highlighted in darker colors. **Left**: uninduced cells, **right**: differentiated cells. **(C)** Volcano plots displaying differential gene expression results based on the BRB-seq^20^ data of expanded SVF-adherent cells from the OM adipose depot versus SVF-adherent cells from other depots (SC, PR, and MC). The top *IGFBP2*+ cell markers identified using scRNA-seq datasets are highlighted in blue, while significantly differentially expressed genes (log2FC > 1, adjusted *p*-value < 0.01) are highlighted in darker colors. **Left**: uninduced cells, **right**: differentiated cells. **(D)** Box plot displaying the log normalized expression of *MSLN* (**top**) and *UPK3B* (**bottom**) across hASPCs (ASCs, PreAs, *HHIP*+, *IFIT*+, *SFRP4*+, *RBP5*+ ASPCs), *IGFBP2*+ cells, Mesothelial cells (Meso) and Vascular Smooth Muscle progenitors (VSMPs), grouped by the depot of origin, as indicated on the x-axis. **(E)** t-SNE cell map of integrated scRNA-seq datasets^8, 12, 13^ from mouse visceral and SC fat depots, depicting the identified clusters: adipose stem cells (ASCs), pre-adipocytes (PreAs), Aregs, *Ifit*+ ASPCs, *Cilp*+ ASPCs, mesothelial, endothelial, and immune cells. **(F)** Box plots showing for each human cell population identified in Figure 2C the score of orthologous murine markers in each mouse cell population as defined in Ferrero et al.^11^.

We classified the first and major population retrieved in all depots as VSMPs, since it expressed muscle-related markers such as *MYH11* but also *ACTA2* and *TAGLN* (**Figures 3A**, and **2E**), resembling a VSMP transcriptomic signature that has previously been described^30^. Noteworthy, beiging of mature adipocytes is accompanied by a shift toward a muscle-like gene expression signature^31–35^, which is why VSMPs may also be involved in thermogenic regulation.

Among the top differentially expressed genes of the ubiquitous *HHIP*+ cluster, we recognized several ortholog markers of a mouse stromal subpopulation that we have previously characterized as having non- and anti-adipogenic properties, and accordingly named Adipogenesis Regulators (Aregs)^12, 17^. These are *F3*, *CLEC11A*, *GDF10*, *MGP*, and *INMT* (**Figure 3A**, **S3A**). Recently, in their single cell atlas of human AT, Emont and colleagues followed by Massier and colleagues identified a cluster that is characterized by enriched expression of *EPHA3*, and that exhibits substantial similarities to the murine Aregs^7, 8^. Notably, *EPHA3* is specifically expressed by the *HHIP*+ cells that we identified in our analyses (**Figure S3A**), further supporting its alignment with mouse Aregs. To solidify the point that the previously described *EPHA3*+ hASPCs are similar to our *HHIP*+ cluster, we transferred our cell annotation onto the Emont et al. dataset^8^ and found that the *EPHA3*+ population has a significantly higher prediction score for our *HHIP+* population than the rest of the hASPCs (**Figure S3B**). Finally, given that *HHIP* is coding for a surface marker, we could confirm the existence of a human SVF Lin–/HHIP+ cell population in the SC AT using flow cytometry (**Figure S3C**-**D**).

Another small stromal population, the *IFIT*+ cluster, which we observed to be present in every depot and donor, is defined by an extremely specific expression of interferon-related genes such as *IFIT3*, *IFI6*, and *IFI27* (**Figure 3A**, **Figure S3E**), a gene signature that is reflective of a viral immune response (**Figure S3F**). A mesothelial *Ifit*+ population has already been reported in mouse OM^15^; yet, our *IFIT*+ population does not express mesothelial markers but mesenchymal ones (**Figure 3A**, **Figure S3G**). However, we found that, based on the expression of ortholog genes between mice and humans, this population shares a very similar signature with *Ifit+* cells that emerged when we integrated multiple mouse ASPC scRNA-seq datasets^11^ (**Figure S3H**).

The *SFRP4*+ cluster was characterized by high expression of Secreted frizzled-related proteins 2 and 4 (*SFRP2* and *SFRP4*) (**Figures 3A**, **S3I**), and aligned with a subpopulation of the published human AT atlas^7, 8^ (**Figure S3J**). SFRPs are inhibitors of the Wnt signaling pathway, a key regulator of adipocyte differentiation^36^, and SFRP2-4, in particular, were shown to be upregulated in obesity, especially in visceral WAT^37^. While the *SFRP4+* population was present in all depots, we observed a general higher expression of *SFRP2*, but not *SFRP4*, in hASPCs from OM adipose depots (**Figure S3K**-**L**).

While the above-described hASPC subpopulations seem to exist in all analyzed adipose depots, albeit at different proportions (**Figure 2D**), we also found three depot-specific cell clusters: the *FMO2*+ cells were specific to PR and MC, and the Mesothelial and *IGFBP2*+ cells to the OM AT (**Figures 2C-D**). The mesothelial cells, defined by the expression of *MSLN, UP3KB, LRRN4*, and Keratin-related genes (**Figure 3A**) constituted an abundant cell type that we retrieved exclusively from the SVF of the OM AT (**Figure 2A, D**). This is consistent with our observation that many Keratin-related genes such as *KRT8*, *KRT9*, *KRT18*, and also *LRRN4* or *UPK1B* were among the top differentially expressed genes in OM cells *versus* those from other depots both at the undifferentiated and differentiated states at the bulk transcriptomic level (**Figure 3B**), which could also explain why the canonical mesenchymal marker *THY1* was less abundant in cultured OM SVF cells, compared to other depots (**Figure S1N**). Similarly, an enrichment of *IGFBP2+* cell markers, including *IGFBP2*, but also others such as *APOE* and *C7* (**Figure 3A**) was also observed in our bulk transcriptomic datasets of OM samples compared to other depots, both at the undifferentiated and differentiated states (**Figure 3C**), thus confirming their specificity to OM. Moreover, when projecting our annotation onto the dataset by Emont and colleagues^8^, our *IGFBP2*+ cluster aligned with one of their clusters (hASPC6) (**Figure 2G** and **S2G**). As a side note, some cells originating from MC samples were also expressing mesothelial markers (**Figure 3D**), in line with the MC AT being itself covered by the peritoneum.

Finally, we systematically mapped each cluster expression score computed on the integrated human scRNAseq dataset (**Figure 2E**) onto the clusters that we have previously identified in mouse^11^ (**Figure 3E-F**) and found high concordance between the proposed nomenclatures. This was further supported by flipping the analysis around and mapping murine cluster expression scores onto the human integrated dataset (**Figure S3M**).

In conclusion, by performing to our knowledge the most comprehensive cross-anatomical analysis of AT-derived stromal cells at the single-cell level, we found five populations that are present in all analyzed depots: the two canonical hASPC subpopulations described before, as well as VSMPs, retrieved in relative high abundance, together with three less abundant stromal populations – *HHIP*+, *IFIT*+ and *SFRP4*+ cells. Specific to the OM SVF were the highly abundant mesothelial cell population and a less abundant *IGFBP2*+ cell cluster. Furthermore, we found high scRNA-seq cluster concordance across the human and mouse models.

### Establishment of a SVF Lin– subpopulation isolation strategy reveals clear phenotypic differences among ASCs, PreAs, and VSMPs

After having characterized the heterogeneity of the cellular SVF Lin– landscape across depots, we aimed at refining our functional characterization between depots at the subpopulation level. We thereby first focused on the main cell populations that are ubiquitous across depots: the ASCs, the PreAs and the VSMPs (**Figure 2B-D**). Based on our scRNA-seq expression profiles, we developed a specific sorting strategy (**Figure 4A**) that would allow the isolation and characterization of each of the aforementioned main SVF Lin– populations. Three layers compose the sorting strategy: 1) the first layer involves CD26, encoded by the gene *DPP4* and specifically expressed by ASCs (**Figure S4A**). Consistent with previous studies^9, 10, 12^, *Dpp4* expression is specific to the murine ASC cluster^11^. 2) The second layer involves Vascular-adhesion protein 1 (VAP1), encoded by the gene *AOC3,* which is highly expressed in VSMPs (**Figure S4A**). In mouse, *Aoc3* expression has mainly been described as being enriched in the PreA population^9, 11, 12^. However, based on our scRNA-seq integration of murine data, *Aoc3* is in fact also highly expressed in murine VSMPs (**Figure S4B**). 3) The third layer aims to enrich for PreAs. Several candidate surface markers appear specific to the PreA population (i.e., *GPC3* or *ICAM1*), however, we reasoned that a simpler PreA enrichment approach would be to select for low expression of CD26 and VAP1. This approach would hold true in every depot except for the OM adipose depot, where two additional OM-specific cell populations would first need to be excluded: the mesothelial and the *IGFBP2*+ cells. Based on our transcriptional analyses, we selected the transmembrane 4 L6 family member 1 (*TM4SF1*) as a marker to first exclude OM-specific populations from downstream functional assays (**Figure S4A** and **C**). In sum, our sorting strategy involves antibodies directed against CD26, VAP1, and TM4SF1 (see **Methods**) to enrich for human ASCs (SVF Lin–/TM4SF1–/CD26+, later referred to as CD26+) and VSMPs (Lin–/TM4SF1–/VAP1+, later referred to as VAP1+), which leaves SVF Lin–/TM4SF1–/VAP1–/CD26– cells, later referred to as DN for “double negative” enriching for PreAs (**Figure 4A-B**).

**Figure 4.**
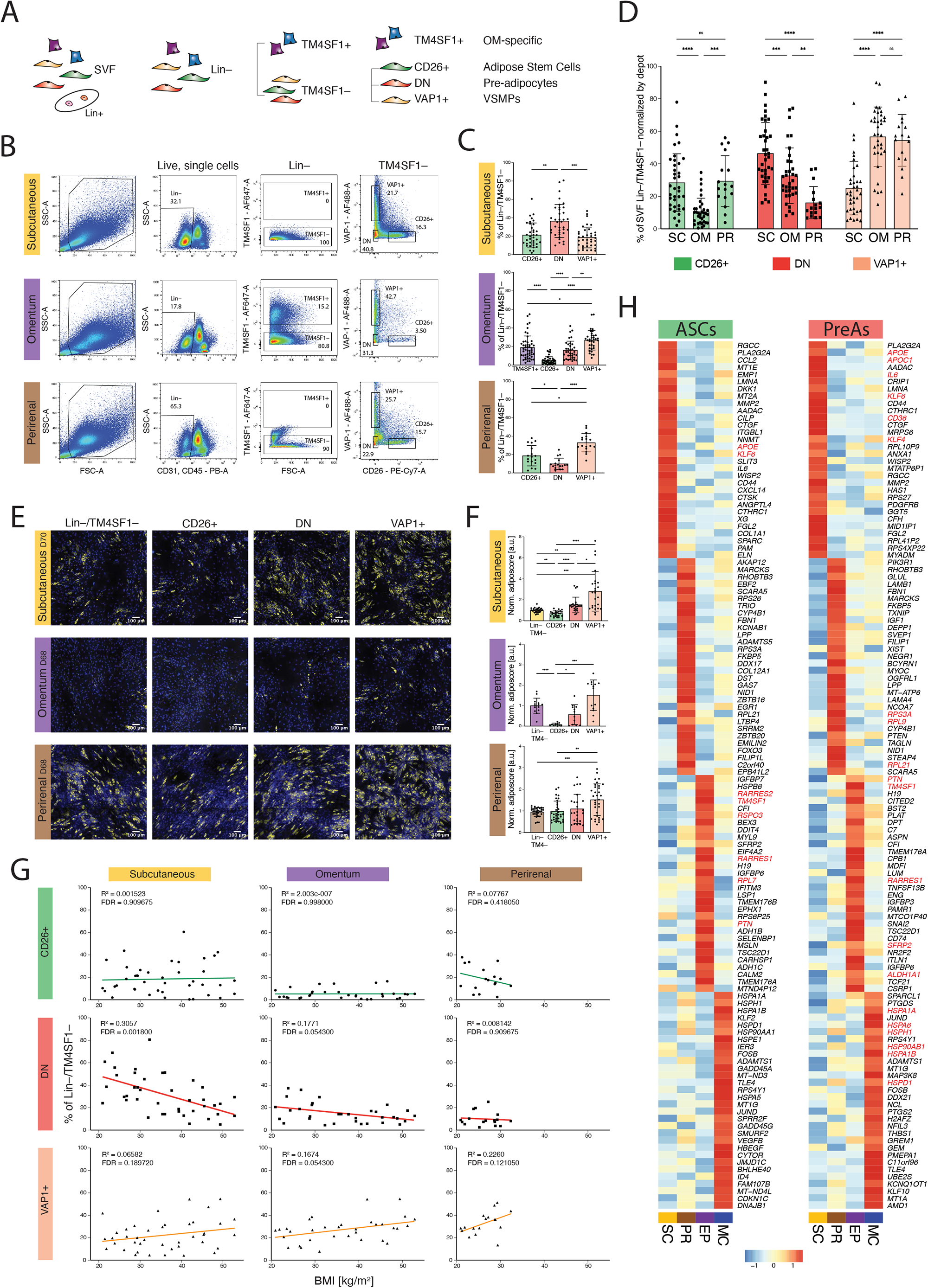
Establishment of a SVF Lin– subpopulation isolation strategy reveals clear phenotypic differences among ASCs, PreAs, and VSMPs. **(A)** Scheme of the sorting strategy used to enrich for adipose stem cells (ASCs), pre-adipocytes (PreAs), vascular smooth muscle progenitors (VSMPs), and omentum (OM)-specific cells. **(B)** Flow cytometry profiles and gating strategy for subcutaneous (SC), OM, and perirenal (PR) SVFs from the same donor (D23) to isolate SVF Lin–/TM4SF1– cells. **(C)** Flow cytometry analysis of the abundance of each cell subpopulation gated from the Lin–/TM4SF1– fraction of SVF cells; SC n=37, OM n=35, PR n=17 donors. **(D)** Bar plot to compare the relative abundance of the indicated SVF populations across depots. The three populations accumulate to 100% of Lin–/TM4SF1– gated cells by depot; SC n=37, OM n=35, PR n=17 donors. **(E)** Representative fluorescence microscopy images of SVF Lin–/TM4SF1–, CD26+, DN, and VAP1+ SVF populations from each depot after *in vitro* adipogenic differentiation (see **Methods**); Yellow – Bodipy staining for lipids, blue – Hoechst staining for DNA, scale bar=100μm. **(F)** Quantification of the adipogenic potential of the SVF Lin–/TM4SF1–populations shown in **E**; Values are normalized to average adiposcore of the reference Lin–/TM4SF1– population; n=12-21, 3-7 donors, 1-4 independent wells each. **(G)** Scatter plot showing the correlation between the % Lin–/TM4SF1– cells from each indicated SVF population and BMI across donors. **(H)** Heatmap of the top 30 higher expressed genes in the indicated depot *versus* all other depots (only genes detected as differentially expressed in each pairwise comparison were retained), focusing on ASCs (**left**) or PreAs (**right**); Average log normalized expression scaled by row. \**p* ≤ 0.05, \*\**p* ≤ 0.01, \*\*\**p* ≤ 0.001, \*\*\*\**p* ≤ 0.0001, One-Way ANOVA and Tukey HSD *post hoc* test (**C**, **D**, **F**), and linear regression analysis with its relative goodness of fit, and the FDR-adjusted *p*-values of the Pearson correlations (**G**).

As expected, and in line with the transcriptomic findings, only OM-derived SVF showed a clearly positive population when stained with anti-TM4SF1 antibody, confirming the exhaustive presence of mesothelial cells in the OM depot (**Figures 4B**, **S4D**). However, as in the scRNA-seq datasets, we did find a few TM4SF1+ cells among MC SVF Lin– cells as well (**Figure S4D**). Analysis of the flow cytometry profiles gathered from up to 37 human donors (**Supp. Table 1**) allowed us to quantify the relative abundance of the targeted populations in each of the three adipose depots (**Figure 4C**). We found that the ASC pool is less abundant in OM AT compared to that of PR and SC, while SC AT is dominated by PreAs and the OM and PR ones by VSMPs (**Figure 4D**). In line with our scRNA-seq findings, we found the same three populations in the MC AT from two donors with relative ratios that resemble those of OM AT (**Figure S4E**-**F**).

Having confirmed the existence of these shared SVF Lin– subpopulations in each depot, we aimed to interrogate their phenotypic behavior *in vitro*. When sorted separately, the CD26+ population outpaced all other populations in terms of cell growth regardless of the depot of origin (**Figure S4G**), a feature that confirms their stem-like nature and is consistent with previous observations in mouse and human^9, 38^. The highly proliferative CD26+ cells also scored the lowest in terms of adipogenic potential (**Figure 4E-F**), further supporting the hypothesis that they are located at the very root of the adipogenic lineage. The VAP1+ cells had the highest adipogenic potential, followed by DN cells (**Figure 4E-F**).

Taking advantage of the cohort of human donors (n=37, **Supp. Table 1**) from which we sampled ATs, we investigated potential correlations between the relative abundance of each of the SVF Lin– subpopulations and corresponding metadata such as BMI, age, and gender of the donors. Interestingly, we found that while the proportion of CD26+ cells (enriching for ASCs) is not affected by BMI changes, the latter appears to be correlated with DN (i.e., PreA) depletion. This anti-correlation is particularly high in the SC, but also in the OM AT and is accompanied by a slight increase in the proportion of VAP1+ cells (enriching for the VSMPs) (**Figure 4G**). In contrast, the age or sex of the donor did not seem to affect the equilibrium of cell populations within the SVF Lin– pool of any of the three analyzed adipose depots (data not shown).

Despite similarities in the transcriptomes of ASCs and PreAs across depots in the scRNA-seq data, we observed that all three OM populations are consistently and significantly less adipogenic than equivalent SC and PR cells. To determine if cell-intrinsic features could explain the low adipogenic capacities of the OM cells, we explored the depot-specific transcriptomic signatures of these subpopulations in our scRNA-seq dataset. We noticed that across depots, the transcriptomes of ASC cells are more related than the PreA ones (**Figure 2G** and **S4H**), supporting the hypothesis that depot-specific features accumulate along commitment. We then identified genes of ASCs or PreAs enriched in a depot-specific manner (**Figure 4H**). In line with their high adipogenic potential, hASPCs from SC, and especially PreAs, showed significantly higher expression of well-known adipogenic genes and transcription factors such *KLF4*, *KLF6*, *WISP2*, *APOE*, *APOC1*, and *CD36*. The pro-adipogenic character of PR-isolated cells was also reflected in their transcriptome (**Figure S4I**). For example, *PIK3R1* is the most up-regulated gene in PR compared to other adipose depots, with PI3K/Akt signaling playing a crucial role in adipogenesis of human mesenchymal stem cells^39^. In mice, PI3K/Akt signaling has also been linked to browning AT by regulating GDF5-induced *Smad5* phosphorylation^40^. It is in this regard of interest that in our scRNA-seq data, *SMAD5* expression was specific to PR PreAs and ASCs. Similarly, *ZBTB16* is a PR-specific marker known to induce browning^41^. With respect to populations that showed limited adipogenic potential, MC cells overexpressed genes linked to unfolded protein or protein folding (**Figure S4I**) such as Heat-shock-proteins (HSPs) (**Figure 4H**), a large family of molecular chaperones. HSPs have been reported to interact with PPARγ to either stabilize it and enhance adipogenesis (Hsp90)^42^ or to destabilize it and inhibit adipogenesis (Hsp20)^43^. OM cells once again showed an enrichment of genes linked to the inflammatory response (**Figure S4I**). Among the candidates specific to OM were also a number of markers that were previously described as having a negative impact on adipogenesis (**Figure 4H**, *RARRES2, RSPO3, RPL7, PTN, GAL, ALDH1A1*, *IGFBP3*^22, 44–46)^.

Taken together, we showed that the hASPC niche harbors different subpopulation abundances depending on the anatomic origin, and its equilibrium changes with increasing BMI. Furthermore, even if ubiquitous across depots, ASCs and PreAs harbor depot-specific gene signatures, seemingly acquired along commitment and potentially reflective of intrinsic phenotypes.

### OM-specific cells inhibit adipogenesis of omental and subcutaneous hASPCs

We next questioned whether the presence of OM-specific cell populations (**Figure 5A**) might influence the adipogenic capacity of the precursor cells themselves, as triggered by two key observations: 1) OM VAP1+ and DN cells, which are depleted of TM4SF1+ cells via the utilized sorting strategy, did show a modest ability to differentiate (**Figure 4E-F**); 2) several genes that were previously linked to the non-adipogenic phenotype of OM SVF-adherent cells were specific to mesothelial and/or *IGFBP2*+ cells (e.g., *CD200*^47^, *WT1*, and *ALDH1A2*^22^, **Figure 5B**).

**Figure 5.**
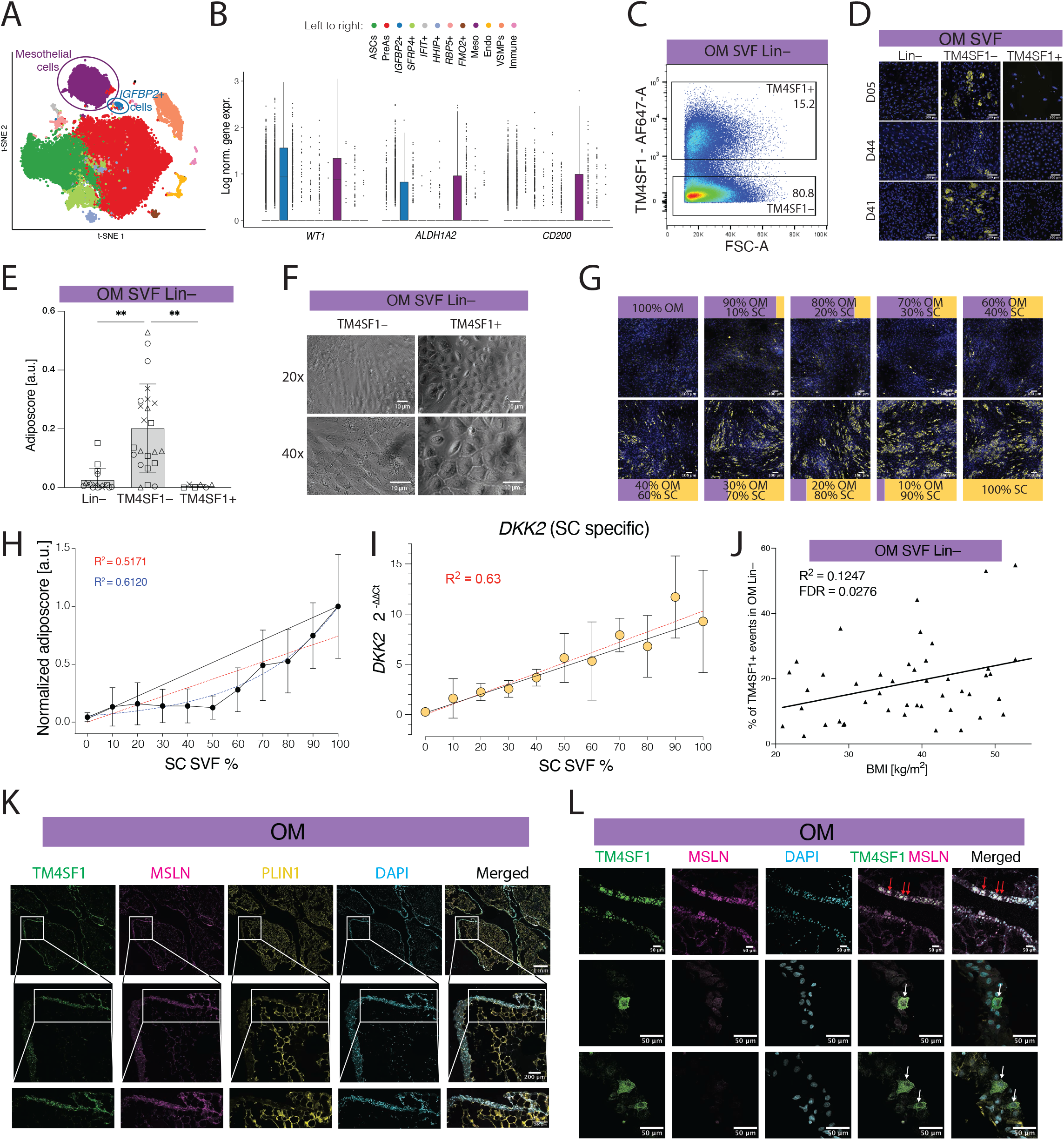
OM-specific cells inhibit adipogenesis of omental and subcutaneous hASPCs. **(A)** t-SNE cell map of integrated scRNA-seq datasets highlighting the two Omentum (OM)-specific populations: Mesothelial cells in purple and *IGFBP2*+ cells in blue. **(B)** Boxplot showing the distribution of log normalized expression of *WT1*, *ALDH1A2*, and *CD200* (x-axis) across the indicated cell populations (defined by the colors), based on the scRNA-seq data in **A**. **(C)** Representative flow cytometry scatter plot of OM SVF Lin– cells (D05) stained with TM4SF1 antibody showing the gating strategy for sorting OM SVF Lin–-specific subpopulations as Lin–/TM4SF1+ and Lin–/TM4SF1– cells. **(D)** Representative fluorescence microscopy images of OM SVF Lin–, Lin–/TM4SF1– and Lin–/TM4SF1+ cell populations after adipogenic differentiation (see **Methods**); Yellow – Bodipy staining for lipids, blue – Hoechst staining for DNA; Scale bars=100μm. **(E)** Barplot showing the adiposcore of the cell populations in **D**; n=6-23, 4 donors, 1-6 independent wells for each. **(F)** Bright-field transmission light microscopy images of spindle-like OM ASPCs (OM SVF/Lin–/TM4SF1–) and cobblestone-like OM-specific TM4SF1+ populations. **(G)** Representative fluorescence microscopy images of SVF Lin– cells in mixing experiments after 14 days of adipogenic differentiation, where SVF Lin– cells from OM and SC adipose tissues of donor 68 (D68) were mixed directly after cell isolation at the indicated proportions. Yellow – Bodipy staining for lipids, blue – Hoechst staining for DNA, scale bar=100μm. **(H)** Adiposcore of the distinct, mixed OM and SC SVF Lin– cell populations, as presented in **G**. Values across biological replicates are normalized to the average adiposcore of the reference 100% SC Lin– condition. The relative proportion (0-100%) of SC SVF Lin– cells in each well is plotted on the x-axis. Error bars represent standard deviation from the average, the linear and exponential regression with corresponding R^2^ coefficients are shown in red and blue, respectively. The black line represents the expected increase of adipogenesis for a linear dilution between 0 and 100% of SC SVF Lin– cells; n=16, 4 biological replicates, 4 independent wells for each. **(I)** qPCR-based gene expression levels of *DKK2* (a subcutaneous depot-specific gene), normalized by *HPRT1* expression and 0% subcutaneous (SC) to control for correct mixing ratios in the experiment shown in **G**. The linear regression and corresponding R^2^ coefficient values are shown in red; a black line links the lowest value to the highest value; n=4, 2 biological replicates, 2 independent wells for each. **(J)** Scatter plot showing the correlation between the OM SVF Lin–/TM4SF1+ fraction based on flow cytometry analysis and the BMI of donors; the line represents a linear regression analysis with its relative goodness of fit; the p-value was computed performing a Pearson correlation. **(K)** Confocal microscopy fluorescent images of the *in situ* immunohistochemistry-based localization TM4SF1 (green), Perilipin (PLIN1) (yellow), and MSLN (pink) cells in OM adipose tissue in donor 67. DAPI staining for nuclei is colored in cyan. These are representative images from 3 independent experiments. **(L)** Confocal microscopy fluorescent images of the *in situ* immunohistochemistry-based localization of TM4SF1+ (green) and MSLN+ (pink) cells in OM adipose tissue in donor 67. DAPI staining for nuclei is colored in cyan. The arrows indicate TM4SF1+/MSLN– cells (white) and TM4SF1+/MSLN+ cells (red) in the periphery of the adipose tissue lobules. Scale bars=50μm. Experiments were repeated at least three times.

Using TM4SF1 as a surface marker for the two OM-specific populations (**Figure S4C**), we depleted the total OM SVF Lin– fraction of TM4SF1+ cells to study the adipogenic behavior of “pure” OM hASPCs (**Figure 5C**). In line with our previous observation on the adipogenic potential of OM DN and VAP1+ subpopulations (**Figure 4E-F**), we found that OM SVF Lin–/TM4SF1– cells, later referred to as OM hASPCs, are significantly more adipogenic than the total OM SVF Lin– fraction. Not surprisingly, since mesothelial cells have previously been shown to be non-adipogenic^48^, the OM SVF/Lin–/TM4SF1+ cells, here referred to as TM4SF1+ cells, did not accumulate lipid droplets (**Figure 5D-E**). This is consistent with their morphological appearance because TM4SF1+ cells stood out from regular spindle-like OM hASPCs^49, 50^ (**Figure 5F**), since they had a round and cobblestone-like shape that is characteristic of mesothelial cells. Importantly, however, the increase in differentiation observed for TM4SF1– cells compared to the Lin– fraction was greater than expected by the simple, proportional removal of the non-adipogenic TM4SF1+ cells (accounting for roughly 20% of the total SVF Lin– fraction, **Figure 4C**). This might suggest that *in vitro* cultured OM hASPCs are subjected to inhibitory cues from the OM-specific TM4SF1+ populations.

To test whether the observed inhibitory cues within the OM SVF Lin– cell pool have a negative influence not only on the adipogenic potential of OM hASPCs but also on those of SC or PR, we set up a mixing experiment where SC Lin– or PR Lin– cells were co-cultured with increasing ratios of OM Lin– cells (**Figures 5G-H** for SC and **S5A-B** for PR). We observed that despite a linear decrease in the relative proportion of OM SVF Lin– cells among SC SVF Lin– ones, the observed increase in adipogenic potential was non-linear (**Figure 5H**). In other words, the increase in differentiation was smaller than expected by the relative proportion of SC SVF Lin– cells. To control for the fact that SC cells were not overgrown by OM cells, we measured the expression of an SC-specific marker, *DKK2* (**Figure S5C**), which revealed no overgrowth as *DKK2* expression showed a linear increase with the proportion of SC cells (**Figure 5I**). Using a similar approach, but this time mixing OM SVF Lin– cells with PR SVF Lin– ones did not reveal any regulatory effect, as we observed a relatively linear relationship between the increase in differentiation and the proportion of PR cells per well (**Figures S5A**-**B**). Thus, our findings suggest that the presence of OM TM4SF1+ cells lowers the adipogenic capacity of neighboring cells, although this effect is not universal among hASPCs and hints at depot-specific sensitivities to the inhibitory cues stemming from OM SVF Lin– cells.

The unexpected ability of OM TM4SF1+ cells to inhibit adipogenesis suggests a possible functional role of this subpopulation in OM AT expansion. This hypothesis is further strengthened by our observation that the relative fraction of OM TM4SF1+ cells within the total SVF Lin– cell pool positively correlated with the BMI of donors (**Figure 5J**). We hence used our scRNA-seq data to resolve this cell population in a more fine-grained manner. This revealed, consistent with results already detailed above (**Figure 2C**), that TM4SF1+ OM-specific cells could be further stratified into two populations: the mesothelial cells and a smaller *IGFBP2*-expressing cluster (**Figures 5A**). To clarify whether the observed inhibition of OM SVF cells over SC SVF cells is specific to one of these two populations, especially given that IGFBP2 itself had previously been described as anti-adipogenic^51, 52^, we aimed at defining an experimental approach to distinguish the two OM-specific populations. To do so, we took advantage of a combination of OM-specific surface markers: 1) we retained TM4SF1 as a marker to enrich for both OM-specific populations together and 2) added MSLN as a marker that is exclusively expressed by mesothelial cells (**Figure S4C**). Hence, we defined mesothelial cells as TM4SF1+/MSLN+ and *IGFBP2*+ cells as TM4SF1+/MSLN– and set out to localize both cell types *in situ* to first validate their *in vivo* presence. The absence of background staining was assessed by both unstained control and secondary-only staining (**Figure S5D**). Interestingly, antibodies directed against both MSLN and TM4SF1 highly stained the boundaries of the AT lobules (**Figure 5K**), likely revealing the mesothelial mono-layer peritoneum structure that pads the OM itself. The majority of positively stained cells were equally intense for both markers; and we defined them as mesothelial cells (**Figure 5L** and **S5D**, red arrows). However, intermingled among these mesothelial cells, we also identified cells that were much more intense in the TM4SF1 channel than the MSLN one (**Figure 5L** and **S5D,** white arrows), reminiscent of our *IGFBP2*+ cell type.

### Omental IGFBP2+ stromal cells appear to transition between mesothelial and mesenchymal cell types

In our scRNA-seq dataset, the *IGFBP2*+ cluster appeared to have an intriguing dual gene expression signature, sharing markers with both hASPCs and mesothelial cells (**Figure 6A**). Such expression signature may at first glance suggest a technical artifact known as doublets, when two cells are mistakenly co-captured and considered as a single one. However, *IGFBP2*+ cells did not display a larger library size or number of captured features (**Figure S6A**), which would be expected for doublets due to a larger initial RNA content compared to singlets. More importantly, we found that these cells express, on the one hand, specific markers such as *IGFBP2*, *RBP1*, *WNT4*, or *WNT6* and, on the other, markers to a higher level than in ASPCs or mesothelial cells alone (**Figure 6B**), which is technically impossible for randomly co-encapsulated cells. To validate the existence of this population in another independent dataset, we transferred our cell annotation onto the recently published snRNA-seq atlas of human SC and OM ATs^8^. We found that, first, only cells from OM harbor a positive prediction score for *IGFBP2*+ cells (**Figure S6B**), validating once more their specificity to the OM. Second, the cells predicted as *IGFBP2*+ cells aligned with a cluster that was independently identified by Emont et al.^8^ (**Figures S6C**-**E**, **S2G**) and showed enrichment for *IGFBP2*+ cell markers, as illustrated by the marker-based expression score (**Figures S6E**). Interestingly, the abundance of this population (relative to ASPCs and mesothelial cells) correlated with the BMI of the donors (ρ=0.95, **Figure S6F**). Once again, aside from expressing their own specific markers (**Figure S6G**-**H**), the predicted cells co-expressed mesothelial and ASPC markers (**Figure S6I**) and aligned along a “bridge” between the two cell types. This duality in gene expression could reflect cells that are transitioning from one cell type to another. To computationally test this hypothesis, we performed trajectory inference on OM hASPCs (ASCs, PreAs), *IGFBP2*+ cells, mesothelial cells as well as VSMPs as a negative control. The trajectory was computed using PAGA, as it can identify continuous and disconnected structures in the data^53^. The inferred graph predicted branches connecting ASPCs to mesothelial cells through *IGFBP2*+ cells (**Figure 6C-D**). As positive and negative controls of the validity of the graph structure, ASCs and PreAs were also connected by a robust branch, as previously reported in mouse^9, 11^, while VSMPs were not connected to the main trajectory. When ordering the cells by their pseudotime along the trajectory starting from ASCs (**Figure 6E**), we observed a gradual decrease and increase of hASPC and mesothelial cell markers, respectively, along the connecting branch (**Figure 6F**), as well as an up-regulation of *IGFBP2*+ cell markers during the transition (**Figure 6G**). Altogether, these results indicate that *IGFBP2*+ cells might represent cells that transition between mesothelial and mesenchymal cell types. Accordingly, we found the GO term “epithelial-to-mesenchymal transition” (EMT) to be enriched among the *IGFBP2*+ cells’ differentially expressed genes (**Figure 6H**). In addition to the genes enriched in the GO term, such as Slug (*SNAI2*), we also found several genes that are expressed by the transitioning cells that were previously linked with EMT, such as genes from the Wnt family, Matrix Metallopeptidase (MMPs), ZEB transcription factors, and others^54–56^ (**Figure 6I**). TGF-β signaling, and especially TGF-β1, has also been described as a master regulator of EMT linked to wound healing and fibrosis^57, 58^. In line, we found that *IGFBP2*+ cells have an enriched expression linked to “response to TGF-β”, but not significantly to TGF-β1 in particular (**Figure 6H**). These cells also express genes in relation to epithelial migration and proliferation. Finally, EMT in the peritoneum of mice has been shown to induce the following gene programs: angiogenesis, hypoxia, inflammatory responses, cell cycle markers, and downregulation of adhesion molecules^59^. The corresponding GO terms were all significantly enriched among the *IGFBP2*+ cell markers (**Figure 6H**). Thus, our findings point to the existence of cells that likely transition between mesothelial and mesenchymal cell types, even under “steady-state-like” conditions.

**Figure 6.**
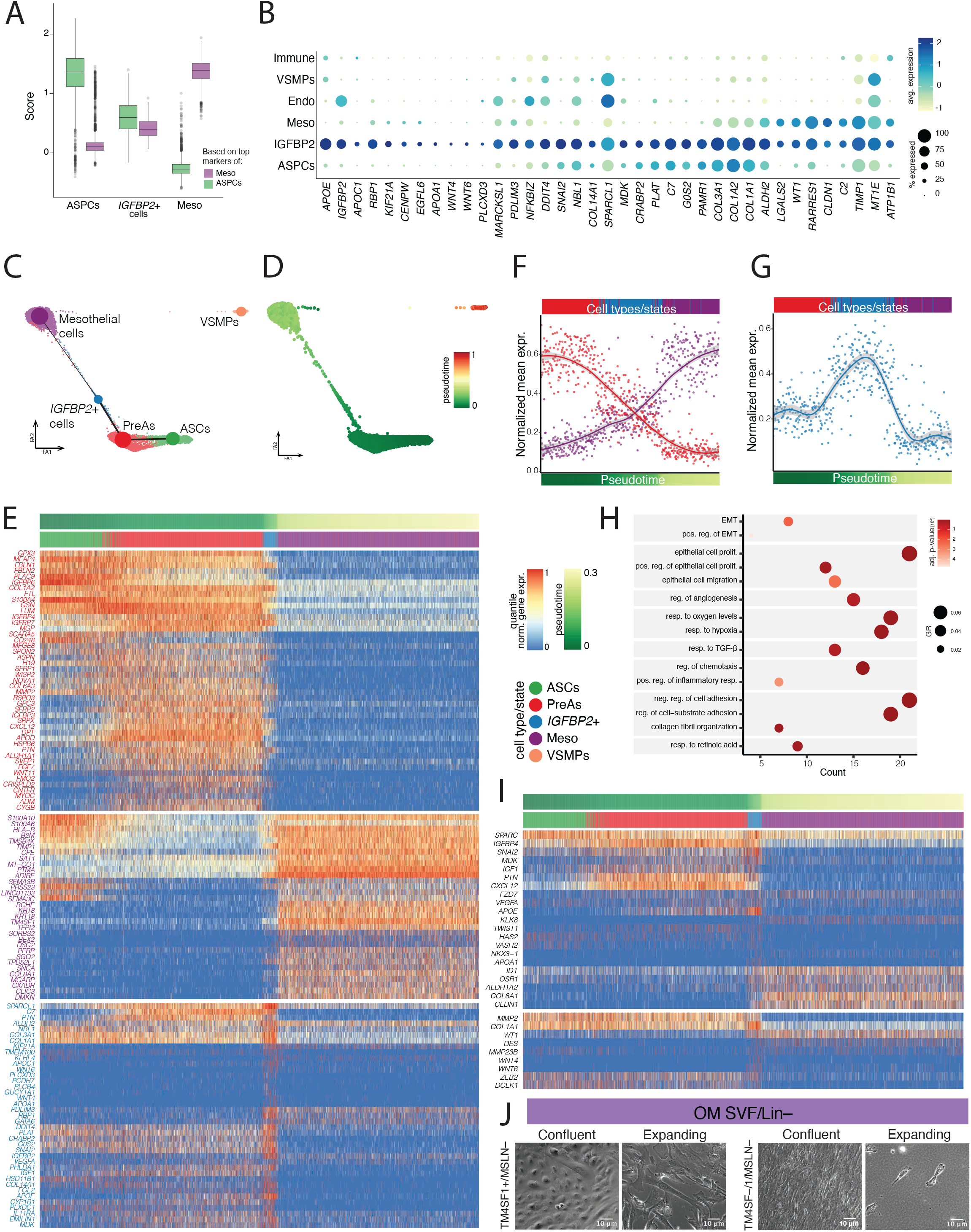
Omental IGFBP2+ stromal cells appear to transition between mesothelial and mesenchymal cell types. **(A)** Box plot showing the distribution of the score based on the top mesothelial cell markers (purple) or the top ASC and PreA markers (green) in OM hASPCs (ASCs and PreAs), IGFBP2+ cells, and mesothelial cells. **(B)** Dot plot displaying the average expression and percentage of expressing cells of the top *IGFBP2*+ cell markers across the clusters shown in Figure 2C. **(C)** PAGA-inferred trajectory superimposed on the PAGA-initialized ForceAtlas2 layout^53^. The size of the dots is proportional to the number of cells in the cluster, and the thickness of the lines is proportional to the confidence of the obtained trajectory relationship. **(D)** PAGA-inferred trajectory described in **C**, colored by the inferred pseudotime (starting from ASCs). **(E)** Heatmap showing the gene expression changes along pseudotime calculated on the trajectory shown in **C**. Genes decreasing from hASPCs (ASCs and PreAs) to Mesothelial cells are highlighted in red, genes increasing from hASPCs to Mesothelial cells are highlighted in purple, and genes specific to *IGFBP2*+ cells are highlighted in blue; log normalized gene expression scaled by row (quantile normalization). **(F)** Scatter plot showing the average of quantile-normalized gene expression highlighted in red or purple on the heatmap shown in **E** for each cell along the pseudotime shown in **D**. The plot focuses on the transition between PreAs (red) and Mesothelial cells (purple), passing by *IGFBP2*+ cells (blue). A locally estimated scatterplot (LOESS) smoothing with 95% confidence interval is shown. **(G)** Scatter plot showing the average of quantile-normalized gene expression highlighted in blue on the heatmap shown in **E** for each cell along the pseudotime shown in **D**. The plot focuses on the transition between PreAs (red) and Mesothelial cells, passing by *IGFBP2*+ cells (blue). A generalized additive model (GAM) fit with 95% confidence interval is shown. **(H)** Dot plot of key GO terms enriched based on *IGFBP2*+ cell markers. **(I)** Heatmap showing the change of gene expression along the trajectory pseudotime shown in **D** for EMT-related genes (top: genes found as enriched when performing GO enrichment analysis, bottom: other EMT-related genes found in the literature). For visualization purposes, the number of cells was downsampled proportionally along pseudotime (see **Methods**). **(J)** Brightfield microscopy images of OM SVF Lin–/TM4SF1+/MSLN– (i.e., IGFBP2+) cells from donor 67 reveal a mesothelial cobblestone-like morphology when confluent and fibroblast spindle-like morphology upon expansion, as opposed to OM SVF Lin–/TM4SF1–/MSLN– (OM ASPCs) cells that display a spindle-like morphology in both situations; Scale bars=10μm.

We pursued our functional validation of this intriguing new cell population by validating a new sorting strategy based on the same combination of markers we used *in situ* (**Figure 5L**). We therefore successfully isolated *IGFBP2*+ cells from the total human OM SVF (see details in the next section). By doing so, and emphasizing their transitioning nature, we found that confluent *IGFBP2*+ cells (OM SVF Lin–/TM4SF1+/MSLN–) harbor the specific mesothelial-cobblestone-like morphology, but when expanding, they tend to adopt a spindle-like shape, resembling mesenchymal cells (OM SVF Lin–/TM4SF1–/MSLN–) (**Figure 6J**).

### Omental IGFBP2+ stromal cells inhibit adipogenesis through IGFBP2

After visualizing cells with low MSLN but high TM4SF1 expression *in situ* by immunohistochemistry (**Figure 5L**), a flow cytometry-based approach allowed us to identify both mesothelial cells (Lin–/TM4SF1+/MSLN+) and IGFBP2+ cells (Lin–/TM4SF1+/MSLN–) *ex vivo* in the SVF of OM biopsies, together with “canonical” OM hASPCs (Lin–/TMSF1–/MSLN–) (**Figures 7A**, **S7A**). To make sure that the gates we set were enriching for our populations of interest, and particularly for the *IGFBP2*+ transitioning cells, we measured *IGFBP2* expression by qPCR in the sorted cells confirming a significant enrichment in Lin–/TM4SF1+/MSLN– cells compared to OM hASPCs and SC SVF Lin– cells (**Figure 7B**). To further validate our sorting and assess whether high *IGFBP2* expression leads to equally high IGFBP2 secretion or intracellular accumulation^60^, we looked for the abundance of the IGFBP2 protein in the supernatant. Using ELISA and concordant to the *IGFBP2* expression measured through scRNA-seq (**Figure S4C**), we measured the concentration of IGFBP2 in the supernatant of confluent OM Lin–/TM4SF1+/MSLN– cells, quantified at approximately 35ng/ml (= 0.97nM). Mesothelial cells secreted less than 20ng/ml (= 0.55nM) of IGFBP2 in similar experimental conditions. In contrast, low IGFBP2 levels were measured in the supernatant of OM SVF Lin– cells, together with barely no IGFBP2 in the supernatants of OM, SC or PR hASPCs (**Figure 7C**). To translate these values to a more physiological model of IGFBP2 secretion by the OM AT, we incubated total OM AT and measured the secreted amount of IGFBP2 after 24, 48, and 72 hours. The concentration of IGFBP2 increased linearly over time, leading to a secretion of ∼5ng/mL for 100 mg of tissue every 24h (**Figure 7D**).

**Figure 7.**
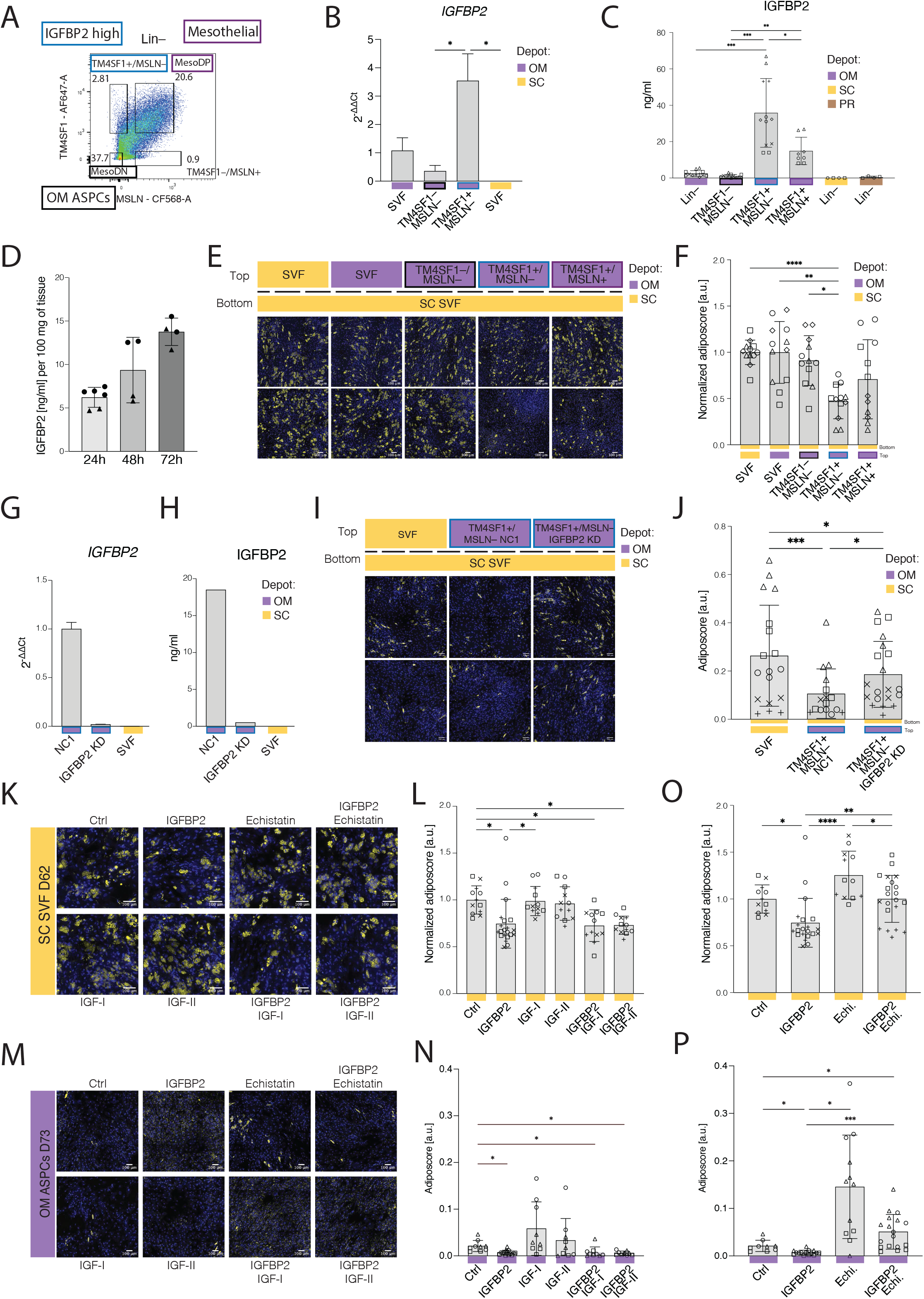
Omental IGFBP2+ stromal cells inhibit adipogenesis through IGFBP2. **(A)** Representative flow cytometry scatter plot of OM SVF Lin– cells (D53) stained with TM4SF1 and MSLN showing the gating strategy to enrich for specific SVF Lin– subpopulations: Lin–/TM4SF1–/MSLN– (OM ASPCs – Black border), Lin–/TM4SF1+/MSLN– (*IGFBP2*+ cells, Blue border), or Lin–/TM4SF1+/MSLN+ (mesothelial cells, Purple border); DP: Double Positive; DN: Double Negative. **(B)** qPCR-based quantification of *IGFBP2* expression. Ct values are normalized first to *HPRT1* expression, then to the ΔCt of OM SVF cells; n=4, 2 donors, 2 technical replicates. **(C)** ELISA-based quantification of secreted IGFBP2 (ng/mL) in the supernatant of the indicated cellular populations after 48h of secretion in a serum-free medium; n=8, 4 donors, 2 technical replicates. **(D)** ELISA-based quantification of IGFBP2 levels (ng/mL), as secreted by 100mg of OM adipose tissue incubated in PBS over the indicated time window; n=4, 2 donors, 2 technical replicates. **(E)** Representative fluorescence microscopy images of “receiver” SC SVF adherent cells, at the bottom of a transwell set-up, after adipogenic differentiation when co-cultured with the indicated SVF populations on top of the transwell: paired SC SVF adherent cells, OM SVF adherent cells, OM SVF/Lin–/TM4SF1– (OM ASPCs), OM SVF/Lin–/TM4SF1+/MSLN– (*IGFBP2*+) cells, or OM SVF/Lin–/TM4SF1+/MSLN+ (mesothelial) cells. Top row: SC cells from D25, OM cells from D54; bottom row: SC and OM cells from D65. **(F)** Bar plot showing the adiposcore quantification of “receiver” cells in **E**. Values are normalized to the average adiposcore of the reference top SC SVF adherent condition; n=12, 4 donors, 3 independent wells. **(G)** qPCR-based quantification of *IGFBP2* expression in SVF/Lin–/TM4SF1+/MSLN– cells subjected to either *IGFBP2* siRNA or non-targeting siRNA control (NC1), as retrieved from the transwell set-up. SC SVF adherent cells are also used as negative control. Ct values are normalized first to *HPRT1* expression, then to the ΔCt of NC1 control; n = 2, 1 donor, two technical replicates. **(H)** ELISA-based quantification of IGFBP2 levels in the supernatant of OM SVF Lin–/TM4SF1+/MSLN– cells subjected to either *IGFBP2* siRNA or non-targeting siRNA control (NC1). SC SVF/Lin– cells are used as negative control; n = 2, 1 donor, two technical replicates. **(I)** Representative fluorescence microscopy images of “receiver” SC SVF adherent cells, at the bottom of the transwell set-up, after adipogenic differentiation when co-cultured, with the indicated cells on top of the transwell: paired SC SVF adherent control cells, OM SVF/Lin–/TM4SF1+/MSLN– cells treated with non-targeting siRNA control (NC1), OM SVF/Lin–/TM4SF1+/MSLN– cells treated with *IGFBP2* siRNA. Top row: SC and OM cells from D74, bottom row: SC cells from D63, and OM cells from D75. **(J)** Bar plot showing the adiposcore quantification of “receiver” cells in **I**; n=16-20, 4 donors, 2-4 independent wells. **(K)** Representative fluorescence microscopy images of SC SVF-adherent cells after adipogenic differentiation when treated with the indicated compounds: IGFBP2 1nM, IGF-I 10nM, IGF-II 10nM, Echistatin 100nM. **(L)** Bar plot showing the adiposcore quantification of cells in **K**, focusing on the IGF-dependent signaling pathway of IGFBP2. Values are normalized to the average adiposcore of the untreated control cells (Ctrl); n=12, 4 donors, three independent wells. **(M)** Representative fluorescence microscopy images of OM SVF/Lin-/TM4SF1-/MSLN-cells after adipogenic differentiation when treated with the indicated compounds: IGFBP2 1nM, IGF-I 10nM, IGF-II 10nM, Echistatin 100nM. **(N)** Bar plot showing the adiposcore quantification of cells in **M,** focusing on the IGF-dependent signaling pathway of IGFBP2. Values are normalized to the average adiposcore of the untreated control cells (Ctrl); n=9, 3 donors, three independent wells. **(O)** Bar plot showing the adiposcore quantification of cells in **K,** focusing on the IGF-independent signaling pathway of IGFBP2. Values are normalized to the average adiposcore of the untreated control cells (Ctrl); n=12, 4 donors, three independent wells. **(P)** Bar plot showing the adiposcore quantification of cells in **M**, focusing on the IGF-independent signaling pathway of IGFBP2. Values are normalized to the average adiposcore of the untreated control cells (Ctrl); n=9, 3 donors, three independent wells. For images in **E**, **I**, **K**, and **M**: Yellow – Bodipy staining for lipids, blue – Hoechst staining for DNA, scale bar=100 um. \**p* ≤ 0.05, \*\**p* ≤ 0.01, \*\*\**p* ≤ 0.001, \*\*\*\**p* ≤ 0.0001, One-Way ANOVA and Tukey HSD *post hoc* test (**B**, **C**, **F**, **L**, **O**), REML analysis with matched values for the same donor and Tukey HSD *post hoc* test (**J**, **N**, **P**).

Given that IGFBP2 is a well-known OM-specific adipokine that has been shown to have anti-adipogenic properties^51, 61, 62^, we wondered if the IGFBP2-secreting cells could exert this effect in a paracrine fashion, accounting for the anti-adipogenic effects of OM over SC cells. To test this hypothesis, we used a transwell setup where receiving cells are exposed to the secretome of either IGFBP2-secreting, mesothelial, or control cells, preventing cell-to-cell contact. At the bottom, we seeded the highly adipogenic SC SVF Lin– cells, and at the top different fractions of OM stromal cells (**Figure 7E**). By doing so, we observed the highest and most significant adipogenic inhibition on SC cells when they were exposed to OM SVF Lin–/TM4SF1+/MSLN– cells, while the adipogenic inhibition was milder and more variable when SC cells were exposed to the OM Lin–/TM4SF1+/MSLN+ fraction (**Figure 7E-F**). To validate that the PR cells are less responsive to this inhibitory signal, as shown in direct co-culture experiments (**Figure S5A**-**B**), we performed the same transwell experiment, but this time with PR SVF Lin– cells at the bottom. Consistent with our first observation, PR hASPCs were rather insensitive to the inhibitory action of OM SVF Lin– cell subpopulations on adipogenesis (**Figure S7B**-**C**).

To directly test whether IGFBP2-secreting cells are inhibitory because of IGFBP2 secretion, we knocked down (KD) *IGFBP2* in the OM SVF Lin–/TM4SF1+/MSLN– cell population using siRNA probes. After validating the KD both at the mRNA and secreted protein levels (**Figure 7G-H**), we used again a transwell set-up to expose SC SVF Lin– cells to the KD cells’ secretome as well as to that of OM SVF Lin–/TM4SF1+/MSLN– cells treated with non-targeting siRNA control (NC1). We found that the SC cells exposed to the *IGFBP2* KD cells were significantly more adipogenic than those exposed to the control IGBFP2-expressing cells (**Figure 7I-J**), further supporting that Lin–/TM4SF1+/MSLN– cells exert an anti-adipogenic action via IGFBP2.

### IGFBP2-mediated adipogenic inhibition occurs in an IGF-independent manner

Prompted by the evidence that IGFBP2 at least partially orchestrates the anti-adipogenic environment observed within OM SVF, we set out to better understand the mechanism underlying IGFBP2’s anti-adipogenic actions. First, we tested if exogenous recombinant IGFBP2 is itself inhibitory by treating SVF-adherent cells from SC or PR depots with increasing IGFBP2 concentrations ranging from 0.25 to 16nM (**Figure S7D**-**E**). We observed that IGFBP2 prevented adipogenic differentiation in a dose-dependent fashion when provided to both SC and PR SVF-adherent cells, albeit remarkedly some PR lines were completely insensitive to the recombinant IGFBP2 treatment. Nevertheless, a significant inhibition of adipogenic differentiation was observed in cells from both depots at concentrations as low as 2nM IGFBP2 (= 72ng/ml). Thus, we used this concentration for the follow-up mechanistic studies (**Figures 7K-P and S7F-H**).

IGFBP2 is known to act through two main mechanisms involving either IGF-dependent or IGF-independent signaling^63^. In the first scenario, the presence of IGFBP2 in the extracellular environment of hASPCs would sequester IGF-I and/or IGF-II and interfere with their pro-adipogenic signaling^64–67^. In the second, IGFBP2 would activate a signaling cascade by binding to the α5β1 integrin receptor, inducing cells to stay in their pre-adipocyte state^67^. Hence, we aimed to narrow down through which of these mechanisms IGFBP2 might influence adipogenesis of hASPCs.

To test whether IGFBP2 acts by sequestering IGFs, we co-treated SVF-adherent cells with both IGFBP2 and IGF-I or IGF-II, as well as with the three recombinant proteins alone. While most literature uses IGF-I and IGF-II at concentrations around 10 nM^65, 67^, we were unable to observe a significant effect on the adipogenic potential of hASPCs treated with IGFs at any concentration ranging from 2.5 to 40nM (**Figure S7D**-**E**). Further, for SC cells, the inhibitory effect of IGFBP2 on adipogenesis was comparable in the presence or in the absence of IGFs (**Figure 7K-L**), suggesting that IGFBP2 influences adipogenesis in an IGF-independent manner. Once again, PR lines appeared to be less sensitive to the action of IGFBP2 and IGF treatments. In fact, even though we observed a similar trend to that observed for SC cell behavior when treating PR cells with IGFBP2 both in the presence or in the absence of IGFs, none of the observed decreases in adipogenic potential were significant when compared to the non-treated cells (**Figure S7F**-**G**). Overall, this is consistent with our previous observations suggesting that PR SVF-adherent cells are less sensitive to the inhibitory effect of OM SVF Lin– cells in the cell mixing setup (**Figure S5A**-**B**) and of OM SVF Lin–/TM4SF1+/MSLN– cells in the transwell setup (**Figure S7B**-**C**).

Next, we explored to what extent OM TM4SF1– cells, enriching for OM hASPCs, can respond to IGFBP2 and IGF treatments, since these cells anatomically co-localize with the IGFBP2-secreting cells. Even if OM TM4SF1– cells are intrinsically lowly adipogenic, we observed an impaired differentiation capacity when these cells were treated with IGFBP2 (**Figure 7M-N**), further supporting the anti-adipogenic capability of IGFBP2-secreting cells in their depot of origin. Contrary to PR and SC cells, OM cells were more sensitive to the IGF-I and IGF-II treatments but with a high degree of variability between batches (**Figure 7M-N**). However, when co-treated with IGFs and IGFBP2, the differentiation of OM TM4SF1– cells was again significantly lower than in non-treated cells (**Figure 7M-N**). The fact that IGF treatment did not influence the actions of IGFBP2 further strengthens the concept of an IGF-independent mode of action by IGFBP2.

We then tested whether IGFBP2 may act in an IGF-independent fashion by activating the α5β1 integrin receptor^68^. To do so, we used echistatin, a known antagonist of the integrin receptor^69^, at a concentration of 100 nM for the first 48h of adipogenic induction^51^, as longer treatment resulted in cell detachment. We therefore coupled echistatin to IGFBP2 treatment only during the first 48h of differentiation. Interestingly, we found that echistatin alone significantly enhanced the differentiation of SC SVF-adherent cells, while, when cells were co-treated with IGFBP2 and echistatin, the adipogenic potential of the treated cells was similar to that of non-treated control cells (**Figure 7K**, **O**). Interfering with integrin receptor function in PR SVF-adherent cells yielded a similar trend in overall adipogenic potential as observed for SC cells (**Figure S7F**, **H**). This result highlights the important role played by integrin receptor signaling in mediating the adipogenic potential of cells, as echistatin had a significant effect even on the highly adipogenic PR cells.

Finally, when treating OM TM4SF1– cells with echistatin, we observed a significant increase in the ability of these intrinsically non-adipogenic cells to accumulate lipid droplets (**Figure 7M**), in line with findings by Yau and colleagues^51^. Furthermore, co-treatment with echistatin and IGFBP2, both competing for binding to the α5β1 integrin receptor, led to a significant increase in differentiation compared to non-treated cells, but less than echistatin-only treatments (**Figure 7M**, **P**).

Taken together, our observations point to the existence of an OM-specific and transitioning cell population that highly expresses and secretes IGFBP2, which negatively impacts the adipogenic potential of OM and SC hASPCs, by signaling through the integrin receptor alpha. However, we cannot completely exclude that the restored adipogenic potential of the analyzed cells (as compared to non-treated control cells) may be driven by two independent and opposite effects, i.e., inhibition by IGFBP2 and enhancement by echistatin. Indeed, the observed significant increase in adipogenesis for example of PR cells upon echistatin treatment (**Figure S7F**, **H**) suggests that the integrin receptor can also negatively regulate adipogenic potential in an IGFBP2-independent manner.

## Discussion

Despite significant efforts, our understanding of hASPC heterogeneity and function across human adipose depots is still limited, in part due to the lack of hASPC consensus markers. To address this, we first performed a comprehensive exploration of human SC, PR, OM, and MC AT SVF Lin– population structure and function. Our bulk analyses revealed extensive molecular and phenotypic variation among these depots (**Figure 1**). On a global level, we confirmed earlier observations that only SVF-adherent cells from extraperitoneal ATs (SC and PR) displayed high adipogenic potential *ex vivo*, while their intraperitoneal counterparts (OM and MC) were refractory to adipogenesis (**Figure 1C**)^19, 70–72^. This is also reflected by the fact that SC and PR SVF-adherent cells featured a highly adipogenic transcriptomic signature compared to OM and MC ones (**Figure 1F** and **S1O**-**Q**), which in contrast featured a more inflammatory and epithelial/mesothelial gene expression profile (OM) ^73^, or a protein trafficking (heat shock protein) expression signature (MC) (**Figure 1J**). However, despite being highly adipogenic, we also found important molecular differences among extraperitoneal ATs, revealing that, contrary to SC, the gene expression profile of PR SVF-adherent cells was enriched for terms associated with the oxidative respiratory chain, thermogenic response, and mitochondrial activity (**Figure 2J**). This suggests that PR hASPCs may be prone to beiging, potentially reflecting an influence of the nearby adrenal gland^72^.

To better explore potential cellular mechanisms underlying the distinct adipogenic properties of the four analyzed depots, we resolved SVF Lin– heterogeneity by performing scRNA-seq on about 34’000 cells (an average of 8’500 cells per depot) and comparing the resulting data with publicly available datasets from both human and mouse ATs^8, 11^. These analyses allowed us to identify stromal populations that are shared across ATs (**Figure 2A-D**), including three relatively small ones, such as *HHIP*+, *IFIT*+ or *SFRP4*+ cells, as well as two main ones: i) the hASCs, which mapped to the mouse *Dpp4*+ population^9, 12, 13^ and the human *DPP4*+ cells^9^, and ii) the hPreAs, which mapped to the mouse *Icam1*+/*Aoc3*+ population^9, 12^ and human *ICAM1*+ clusters^9^. The ASC pool is proportionally the smallest in OM AT (**Figure 4D**), supporting the hypothesis that SC and PR ATs have a greater capacity to expand through hyperplasia compared to OM AT^74, 75^. A third cluster that was ubiquitous in all analyzed human depots is the VSMP cluster which highly expresses *AOC3* (VAP1) (**Figure 2A-D** and **S4A**). Although *Aoc3* has mainly been described as being expressed by murine PreAs^9, 12^, murine VSMPs do exist and also highly express *Aoc3* (**Figure S4B**). As human PreAs also exhibit basal *AOC3* expression, we cannot completely rule out that VAP1 also enriches for a fraction of human *AOC3*-expressing PreAs. In our study, VAP1+ cells were the most adipogenic (**Figure 4E-F**), but at the transcriptomic level, *AOC3*-high cells also expressed muscle-related markers (**Figure 3A**), which seems contradictory. However, beige/brown AT progenitors have been described to upregulate muscle-related markers to become thermogenic^31–35^. Thus, we cannot exclude that VSMP and/or VAP1-enriched PreAs might act as beige progenitors. The fact that VAP1+ cell abundance was high in OM and PR ATs would be in agreement with the observation that, contrary to mice, human visceral AT can also undergo beiging^10, 76–78^. Interestingly, VAP1+ cells showed a greater abundance in high *versus* normal weight individuals across all analyzed adipose depots (**Figure 4G**). This may reflect an attempt to either induce a thermogenic response to balance excessive energy take or to create new vasculature to support adipose tissue expansion.

The above results highlight the many similarities found between human and mouse ASPCs. However, we could also detect some clear differences. For example, while *F3*+ ASPCs form a clearly distinct cluster in mouse visceral and subcutaneous-derived scRNA-seq datasets^9, 11, 12, 17, 18^, they appear to be less abundant in humans (**Figure 2C-D**). Moreover, while *F3* is a specific marker for this anti-adipogenic stromal populations in mice, it is much less specific in humans, where *HHIP* appears to be a more specific marker for this cell population (**Figure 3A**).

In addition to the AT-ubiquitous cell populations, we also identified populations that are specific to one adipose depot. A striking example are the mesothelial cells that are almost exclusive to OM AT (**Figures 2A, D**, **3D**). While the presence of mesothelial cells within the OM SVF has been reported previously^8, 13, 15, 16^, their role within the adipose stem cell niche remained elusive. Our functional characterization revealed that these mesothelial cells can inhibit the differentiation of OM hASPCs (**Figure 5D-E**), suggesting that the mesothelium surrounding the OM AT could have a regulatory impact on its plasticity. Our work suggests that the anti-adipogenic action of omental mesothelial cells is driven by a specific subpopulation that could be sorted as OM SVF/Lin–/TM4SF1+/MSLN– cells. These cells highly secrete IGFBP2 (**Figure 7C**) and strongly repress the adipogenic capacity of both SC and OM hASPCs (**Figures 5D-E**, **7E-F**). This is consistent with IGFBP2’s previously reported anti-adipogenic properties^51, 79^. Mechanistically, our findings revealed that the anti-adipogenic property of Lin–/TM4SF1+/MSLN– cells is modulated by the secretion of IGFBP2 (**Figure 7I-J**) which acts through an IGF-independent mechanism, most likely via the activation of integrin receptor signaling (**Figure 7K-P**). The identification of this cell population might help explaining the limited adipogenic capacity of OM hASPCs in culture. However, the knockdown of IGFBP2 only partially rescued the ability of OM hASPCs to be adipogenic (**Figure 7I-J**). This indicates that OM hASPCs still feature cell-intrinsic and transcriptomically independent mechanisms that render them refractory to differentiation *ex vivo*.

Our identification of an OM-specific anti-adipogenic cell lines evokes the discovery in mice of Aregs, which are stromal populations that negatively regulate the adipogenic capacity of ASPCs in mouse subcutaneous ATs, both by our^12, 17^ and other labs^16, 18^. These discoveries suggest that, also in humans, AT plasticity may be orchestrated by distinct cues including not only endocrine signals but also specialized niche cells. However, classical Aregs and OM-derived IGFBP2-secreting cells have a very different cellular identity. While Aregs are of mesenchymal nature, we found that *IGFBP2*+ cells expressed a joint mesenchymal and mesothelial identity (**Figure 6A**) and showed enrichment of mesothelial to mesenchymal transition (MMT) markers (**Figure 6H-I**). Moreover, when freshly sorted as TM4SF1+/MSLN– cells, they exhibited a cobblestone-mesothelial morphology while, upon expansion, a spindle-mesenchymal one (**Figure 6J**), further suggesting their capacity to undergo MMT, a still poorly characterized process that has been described to also be driven by IGFBP2 itself^80–83^. While this cellular process is known, it has mainly been described in development, wound healing and cancer. Our results suggest however that MMT can also occur in adulthood. Interestingly, by projecting our annotation onto the recently published single-cell atlas of human AT^8^, we made two interesting observations on how *IGFBP2+* cells might relate to human (adipose) biology. First, we found that *IGFBP2*+ cells can be detected in the OM adipose depots of both lean and obese donors (**Figure S6F**). Second, we also observed a highly positive correlation between inferred *IGFBP2*+ cell abundance and BMI (**Figure S6F**). The latter observation appears to contrast with results from previous studies reporting an anti-correlation between BMI^84–86^, onset of metabolic syndrome^87^ including type 2 diabetes and NAFLD^88^ on the one hand and circulating IGFBP2 serum levels on the other. One possible explanation is that a higher number of *IGFBP2*+ cells does not necessarily mean a higher level of expression or secretion. Also, since IGFBP2 is also secreted by other organs such as the liver^61, 88^, additional research is required to reconcile IGFBP2’s paracrine actions controlling local OM AT plasticity versus systemic actions as a metabolic regulator.

Altogether, our work contributes to a better understanding of the behaviors of different human fat depots, some of which are still poorly explored in the literature. It also highlights the main cellular populations that are conserved across depots and species. And, finally, it identifies and mechanistically characterizes an OM-specific population that inhibits the differentiation of neighboring ASPCs. While an important proportion of human visceral fat is contained in the OM, this depot is rather minimal in mouse^89^. It may therefore prove difficult to find an equivalent population in mice. However, a very recent study by Zhang et al.^16^ of mouse epididymal AT did identify “mesothelial-like cells’’ that shared markers with both mesothelial and mesenchymal cells and that were also defined by high *Igfbp2* expression. This suggests that OM *IGFBP2*+ cells may be cellularly and functionally conserved between mouse and human, which in turn may open new experimental avenues to study their relevance in mediating OM AT plasticity in distinct metabolic contexts. A better understanding of the action of OM I*GFBP2+* cells could also lead to new therapeutic strategies to render OM hASPCs more adipogenic and less inflammatory, which could be a valuable novel approach to treat metabolic disorders linked to obesity^86^.

## Methods

### Bioethics

All materials used in this study have been obtained from AT donors from two independent cohorts: the Cohort of Obese Patients of Lausanne with ethically approved license by the commission of the Vaud Canton (CER-VD Project PB_2018-00119) and a control healthy cohort from renal transplantation donors with ethically approved license by the commission of the Vaud Canton (CER-VD 2020-02021). The coded samples were collected undersigned informed consent conforming to the guidelines of the 2000 Helsinki declaration. **Supp. Table 2** illustrates cohorts demographics.

### Human ASPCs isolation and culture

2-3 cm^3^ biopsies from SC, OM, PR and MC ATs were washed in PBS to remove excess blood, weighted and finely minced using scissors. Minced adipose tissue was incubated with 0.28 U/ml of liberase TM (Roche #05401119001) in DPBS with calcium and magnesium (Gibco #14040091) for 60 min at 37 °C under agitation. Vigorous shaking was performed after 45 min of incubation to increase the yield of recovered SVF cells. The digested tissue was mixed with an equal volume of 1% human albumin (CSL Behring) in DPBS −/− (Gibco #14190094) to stop the lysis. Following a 5-min centrifugation at 400 g at room temperature, floating lipids and mature adipocytes were discarded by aspiration and the resuspended SVF pellet was sequentially filtered through 100-μm and 40-μm cell strainers to ensure a single cell preparation. To lyse red blood cells, pelleted SVF was resuspended in VersaLyse solution (Beckman Coulter #A09777) according to the manufacturer’s recommendations and washed once with 1% albumin solution. Obtained red blood cell-free SVF suspension was then either plated for experiments, expanded and cryoprotected or stained for sorting (see below). The SVF used for expansion or experiments was plated at a density of at least 100’000 cells per square centimeter in high glucose MEMalpha GlutaMax medium (Gibco #32561037) supplemented with 5% human platelet lysate (Sigma #SCM152) and 50 μg/ml Primocin (InvivoGen #ant-pm-2). For culturing human ASPCs, TrypLE Select reagent (Gibco #12563011) was used to collect the cells from the cell culture plates.

### Bulk RNA barcoding and sequencing (BRB-seq)

All cells for BRB-seq were seeded in parallel in six 24-well plates. Cells from three wells were harvested undifferentiated (t0 time point) upon cell expansion in the 24-well plate. Cells from the three remaining wells were expanded until confluence and harvested in TRIzol (Sigma, #T3934) after 14 days of adipogenic differentiation (t14 time point). RNA was extracted from all samples in parallel using the Direct-ZOL 96 well plate format (Zymo, #R2054), and BRB-seq libraries were prepared as previously described ^20^ and further detailed by the Mercurius^TM^ Protocol (Alithea Genomics). In brief, 7-200 ng of total RNA from each sample was reverse transcribed in a 96-well plate using SuperScriptTM II Reverse Transcriptase (Lifetech 18064014) with individual barcoded oligo-dT primers, featuring a 12-nt-long sample barcode (IDT). Double-stranded cDNA was generated by second-strand synthesis via the nick translation method using a mix containing 2 μl of RNAse H (NEB, #M0297S), 1 μl of *E. coli* DNA ligase (NEB, #M0205LL), 5 μl of *E. coli* DNA Polymerase (NEB, #M0209LL), 1 μl of dNTP (10 mM), 10 μl of 5x Second Strand Buffer (100LmM Tris, pHL6.9, (AppliChem, #A3452); 25LmM MgCl_2_ (Sigma, #M2670); 450LmM KCl (AppliChem, #A2939); 0.8 mM β-NAD (Sigma, N1511); 60 mM (NH_4_)_2_SO_4_ (Fisher Scientific Acros, #AC20587); and 11 μl of water was added to 20 μl of ExoI-treated first-strand reaction on ice. The reaction was incubated at 16 °C for 2.5 h. Full-length double-stranded cDNA was purified with 30 μl (0.6x) of AMPure XP magnetic beads (Beckman Coulter, #A63881) and eluted in 20 μl of water.

The Illumina-compatible libraries were prepared by tagmentation of 10-40 ng of full-length double-stranded cDNA with 1 µl of in-house produced Tn5 enzyme (11 μM). After tagmentation, the libraries were purified with DNA Clean and Concentrator kit (Zymo Research #D4014) eluted in 20 µl of water and PCR amplified using 25 μl NEB Next High-Fidelity 2x PCR Master Mix (NEB, #M0541LL), 2.5 μl of each i5 and i7 Illumina index adapter (IDT) using the following program: incubation 72 °C—3 min, denaturation 98 °C— 30Ls; 15Lcycles: 98 °C—10 s, 63 °C—30 s, 72 °C—30 s; final elongation at 72 °C— 5Lmin. The libraries were purified twice with AMPure beads (Beckman Coulter, #A63881) at a 0.6x ratio to remove the fragments < 300 nt. The resulting libraries were profiled using a High Sensitivity NGS Fragment Analysis Kit (Advanced Analytical, #DNF-474) and measured using a Qubit dsDNA HS Assay Kit (Invitrogen, #Q32851) prior to pooling and sequencing using the Illumina NextSeq 500 platform using a custom primer and the High Output v2 kit (75Lcycles) (Illumina, #FC-404-2005). The library loading concentration was 2.4 pM, and the sequencing configuration was as follows: R1 21c / index i7 8c / index i5 8 c/ R2 55c.

In parallel, the same cells were seeded in four independent 96well plates and imaged after 14 days of differentiation to quantify their adipogenic potential (see “*In vitro* adipogenic differentiation of hASPCs”).

### Analysis of BRB-seq data

#### Preprocessing

After sequencing and standard Illumina library demultiplexing, the*.fastq* files were aligned to the human reference genome GRCh38 using STAR (Version 2.7.3a), excluding multiple mapped reads. Resulting BAM files were sample-demultiplexed using BRB-seqTools v.1.4 (https://github.com/DeplanckeLab/BRB-seqTools) and the “gene expression x samples” read, and UMI count matrices were generated using HTSeq v0.12.4.

#### General methods

Samples with a too low number of reads or UMIs were filtered out. Genes with a count per million greater than 1 in at least 3 samples were retained. Raw counts were then normalized as log counts per million with a pseudo count of 1, using the function `cpm` from *EdgeR*^90^ version 3.30.3. If the samples were from different batches, the raw counts were first normalized using quantile normalization as implemented in `voom` from the package *limma*^91^ version 3.44.3 and then corrected for batch effects using `combat` from *sva* version 3.36.0. PCAs were computed using `prcomp` with the parameters center and scale set to TRUE. Differential expression analyses were performed using *DESeq2*^92^ version 1.28.1 and adding batch as a cofactor when necessary.

#### Scores

Scores were calculated as the sum of the integrated gene expression scaled between 0 and 1 per gene of the mentioned gene lists.

#### Gene expression heatmaps

Heatmaps display row-normalized expression and were generated using `pheatmap` version 1.0.12. The columns and rows were clustered using the method “ward.2D” of `hclust` of the package *stats*.

#### Gene set enrichment analysis

Gene set enrichment analysis was performed using the package *clusterprofiler*^93^ version 3.16.1.

### scRNA-seq of SVF Lin– cells

SVF Lin-cells from different depots and donors were enriched with either FACS or MACS (**Supp. Table 3**) and resuspended in 1% human albumin in DPBS solution prior to be loaded into the Chromium Single Cell Gene Expression Solution (10x Genomics), following the manufacturer’s recommendations targeting a recovery of 4000 to 5000 cells per run. scRNA-seq libraries were obtained following the 10x Genomics recommended protocol, using the reagents included in the v2 or v3 Chromium Single Cell 3′ Reagent Kit depending on samples (**Supp. Table 3**). Libraries were sequenced on the NextSeq 500 v2 (Illumina) instrument using 150 cycles (18 bp barcode + UMI, and 132-bp transcript 3′ end), obtaining ∼5 × 108 raw reads.

### Analysis of scRNA-seq data

#### Analysis of the datasets individually

Raw fastqs were processed using the default CellRanger pipeline (v 2.1.0, 10X Genomics, Pleasanton, CA). The same transcriptome version was used to align all the datasets (GRCh38.92). All the data were then loaded on R (R version 3.6.1). Cells were filtered for the number of Unique Molecular Identifiers (UMIs) and genes using `isOutlier` from the package *scater*, which determines which values in a numeric vector are outliers based on the median absolute deviation (MAD) (nmads set between 3 and 4), and filters for too high a percentage of UMIs mapping to mitochondrial RNA (∼10%) or ribosomal RNA (∼20%) or too low a percentage of UMIs mapping to protein-coding genes (∼80%).

The datasets were first analyzed one by one using the Seurat pipeline ^94^. After cell filtering, only genes expressed in at least 3 cells were kept. The data were scaled for the number of UMIs and features using the function `ScaleData` and the remaining default parameters. The first 50 principal components of the PCA were computed using `RunPCA`, and then evaluated for significance using the `JackStraw` function of Seurat. Only the first PCs successively having a p-value < 0.05 among the top 50 PCs were selected for downstream analysis. Clustering was performed using `FindNeighbors`. The robustness of the clustering was assessed using `clustree` displaying the relationship between the clusters with increasing resolution. Differential expression analysis was computed using the `FindAllMarkers` function of Seurat for the selected clustering. Only genes detected as differentially expressed (log_2_FC > log_2_(1.2), p.adj < 0.05) for both the Likelihood-ratio test (test.use = “bimod”) and Wilcoxon Rank Sum test (test.use = “wilcox”) were selected.

Each sample was processed and sequenced individually, with the exception of the samples PR – D30 and PR – D61. The isolated cells of these two samples and donors were mixed. The cells were identified as belonging to each donor post-processing based on two criteria: the results of the clustering of the dataset, which clearly separated the cells from the two individuals, and the expression of *XIST* as the two donors were of the opposite sex. Cells ambiguously assigned to a donor (i.e, having a positive expression of *XIST* while clustering with the cells of the donor patient or the opposite) were filtered out.

#### Comparison of top markers of individual datasets

For each pair of subpopulations and dataset, the percentage of shared markers between their top 100 differentially expressed genes with the highest FC were calculated and displayed on **Figure S2A**-**C**.

#### Scmap

The *Scmap* package^95^ was used to project the cells of a dataset X onto the identified subpopulations of a dataset Y. Each pair of dataset X, Y and its inverse Y, X were computed. More precisely, the datasets were normalized using the “Single-cell Analysis Toolkit for Gene Expression Data in R” (*scater* package). The data were log normalized using the `logNormCounts` functions using the size factor estimated with `compteSumFactors`. The 1000 most informative features of each dataset were selected using the `selectFeatures` function of *scmap,* which is based on a modified version of the *M3Drop* method. The centroids of each cluster for each dataset were calculated with the function `indexCluster`, and finally, the datasets were projected onto one another using the function `scmapCluster`.

#### Data integration

The datasets from each individual patient and depot, at the exception of GB-D07 (due to a very low number of captured ASPCs), were integrated following the standard workflow of Seurat pipeline. The datasets were normalized in log scale with a scale factor of 10000. The top 2000 highly variable genes were selected using the `FindVariableFeatures` function with the parameter selection.methods set to “vst”. The anchors were identified using `FindIntegrationAnchors`. The top 2000 variable features identified by `SelectIntegrationFeatures` and the first 60 principal components of the PCA were used as input to perform canonical correlation analysis. The integrated data computed by `IntegrateData` were then used for dimensionality reduction and clustering based on the first 60 principal components of the PCA. Clustering was computed for different clustering resolutions. The final clustering result was based on the clustering results at different resolutions depending on the robustness of the clusters and the specificity of their differentially expressed markers. Top differentially expressed genes were identified using the `FindConservedMarkers` function of Seurat after setting the default assay to RNA, the adjusted p-values were combined using Tippett’s method as implemented by the function `minimump` from *metap* R package (meta.method = metap::minimump)^96^. Only groups of cells with at least 10 cells were tested (min.cells.group = 10). Specifically, for the *IGFBP2*+ cell cluster, as we found only a few cells per batch and we focused on that cell type in part of the manuscript, DEGs were further computed using *EdgeR* and correcting for batch. More precisely, genes not expressed in at least 2% of the cells were filtered out using the function `filterByExpr`. After converting the count matrix into a DGEList using `DGEList`, the data were normalized with `calcNormFactors`. The design matrix was defined following the formula ∼0 + clust + batch, where clust corresponds to the cluster of every cell and batch to its dataset (as individually shown on **Figure 2A**). The dispersion was estimated using `estimateDisp`. The quasi-likelihood negative binomial generalized log-linear model was fitted using `glmQLFit`, followed by the quasi-likelihood *F*-test `glmQLFtest` contrasting the *IGFBP2*+ cluster *versus* the other clusters (pondered by the number of clusters).

#### Identification of depot-specific markers for ASCs and PreAs

DEG analysis was performed on the integrated data, by selecting the cells of the population of interest (ASCs or PreAs) and contrasting between all possible pairs of depots using the function `FindMarkers` of Seurat. This is possible as we have 3 replicates for SC, OM, PR, and 2 for MC, however, for the latter, those were coming from two biological samples from the same donor. A set of markers was considered depot-specific when significantly differentially expressed in a depot *versus* any other depot. A gene was defined as differentially expressed when its average log Fold Change (defined as the average of the log Fold Change in each replicate) was positive and an adjusted p-value smaller than 0.05.

#### Comparison with murine ASPCs

##### a. Murine data integration

The integration of five datasets of adult mouse SC and OM ATs provided by Schwalie et al.^12^, Burl et al.^13^, Hepler et al.^14^ and Merrick et al.^9^ was performed as described in Ferrero et al.^11^. The clustering originally published in Ferrero et al.^11^, focusing on ASPCs, merged the cells close to endothelial cells into one main cluster. The clustering was here revised to include vascular smooth muscle progenitor cells. For consistency with the human data, the top markers of the subpopulation were computed as defined above. The top markers were ordered by the average of the log_2_ Fold Change of each dataset.

##### b. Score

Scores of the mouse ASPC subpopulations, mesothelial cells, and vascular smooth muscle progenitor cells were based on their human orthologs and calculated as the sum of the gene expression scaled between 0 and 1 per gene of the top markers (average log_2_ Fold Change across batches > 0 and adjusted p-value < 0.05) of each murine ASPC subpopulation (ASCs, PreAs, Aregs, *Ifit*+, and *Cilp*+ ASCs), mesothelial cells and vascular smooth muscle progenitor cells. The scores were then scaled by the number of genes on each list.

#### Comparison with the dataset from Emont et al.^8^

The whole human single-nucleus/cell dataset (here reported as “scRNA-seq”) provided by Emont *et al.*^8^ was downloaded on the single cell portal (study no. <u>SCP1376</u>, All cells). The dataset was then subsetted for the cells defined as ASPC or mesothelium by the authors (as defined in the metadata “cell_type2”), and the PCA was recomputed as well as clustering, tSNE and UMAP with the first 50 PCs as input. First, an *IGFBP2* expression score was computed using the AddModuleScore function. The dataset containing only ASPCs, and mesothelial cells was then split by samples, and the symbol gene IDs were converted to Ensembl ID using the GRCh38 release 92 from the Ensembl gene annotation as reference. The few genes with no corresponding Ensembl IDs were filtered out, and, in the rare case of two corresponding Ensembl IDs, only one was kept. Each sample was log normalized with the default normalization of the *Seurat* package and then scaled for the features selected using SelectIntegrationFeatures with each of the samples of Emont et al.^8^ and our generated single-cell SC and OM datasets as input. The first 50 PCs were computed based on the scaled data. Clustering was performed following the default Seurat clustering pipeline for resolutions spanning from 0.1 to 3. Each sample of the Emont et al.^8^ dataset was then projected on our integration (see *Analysis of single-cell RNA-seq, Data integration*), using the FindTransferAnchors and TransferData functions of the *Seurat* package with the default parameters.

#### Trajectory analysis

Trajectory analysis was performed on the integrated normalized data subsetting for Epiploic samples. Potential doublets were excluded from the analysis using `DoubletFinder`^97^ on each epiploic scRNA-seq dataset individually. Cells labeled as ASCs, PreAs, *IGFBP2+* cells, Mesothelial cells, and VSMPs were selected. The first 50 PCs were computed using the `pca` function of *scanpy*^98^ and the neighborhood graph was computed with the default parameters (`pp.neighbors`). The connectivity between our defined cell classifications was computed using the `paga` function^53^, and low-connectivity edges were thresholded at 0.03. We computed the ForceAtlas2 (FA2) graph^99^ using PAGA-initialization (`draw_graph`). The *Dynverse* package^100^ was used to compute the most variable genes along the branch connecting PreAs and Mesothelial cells through *IGFBP2*+ cells (`calculate_branch_feature_importance`).

### FACS sorting of human SVF subpopulations

SVF cells were resuspended in 1% human albumin solution (CSL Behring # B05AA01) in PBS to the concentration of 10^6^ cells/μl, and the staining antibody panels (**Supp. Table 4**) were added in titration-determined quantities. At first, all SC, OM, and PR cells were stained with the OM-specific panel, including mesothelial markers, but since SC and PR SVF cells were consistently negative for the TM4SF1 and MSLN markers over three consecutive experiments, SC and PR cells were only stained with the SC and PR panels, respectively (**Supp. Table 4**). The cells were incubated with the cocktail of antibodies on ice for 30 min protected from light, after which they were washed with 1% human albumin in PBS and stained with propidium iodide (Molecular Probes #P3566) for assessing viability, and subjected to FACS using a Becton Dickinson FACSAria II sorter or a MoFlo Astrios EQ, Cell Sorter – Beckman Coulter. Compensation measurements were performed for single stains using compensation beads (eBiosciences #01-2222-42).

The following gating strategy was applied while sorting non-hematopoietic and non-endothelial cells: first, the cells were selected based on their size and granularity or complexity (side and forward scatter), and then any event that could represent more than one cell was eliminated. Next, the live cells were selected based on propidium iodide negativity, and from those, the Lin– (CD45–/CD31–) population was selected. For the SC samples, from the Lin– fraction of cells, Lin–/CD26+, Lin–/VAP1+, Lin–/DN, and Lin–/HHIP+ cells were defined against unstained controls and FMO controls. For the PR samples, from the Lin– fraction of cells, Lin–/CD26+, Lin–/VAP1+, and Lin–/DN cells were defined against unstained controls and FMO controls. For the OM samples, OM-specific subpopulations were first isolated from the Lin– gate as Lin–/TM4SF1+/MSLN– and Lin–/TM4SF1+/MSLN+ populations. From the remaining Lin–/TM4SF1– gate, we then isolated Lin–/TM4SF1–/CD26+, Lin–/TM4SF1–/VAP1+, and Lin–/TM4SF1–/DN cells. Acquired FCS files were analyzed using FlowJo software to infer population abundances that were plotted using GraphPad Prism.

### *In vitro* adipogenic differentiation and chemical treatments of hASPCs

Cells were seeded for adipogenic differentiation at high density (65k cells /cm^2^) in 3-5 replicate wells of a 96-well black plate (Corning #353219). After 48h or when cells where confluent for at least 24h, cells were treated with induction cocktail (high glucose DMEM (#61965), 10% FBS, 50 μg/ml Primocin, 0.5 mM IBMX (Sigma #15879), 1 μM dexamethasone (Sigma #D2915), 1.7 μM insulin (Sigma #19278), 0.2 mM indomethacin (Sigma #I7378) for 7 days, followed by a maintenance cocktail (high glucose DMEM, 10% FBS, 50 μg/ml Primocin, 1.7 μM insulin) for another 7 days. No medium refreshment was performed between these two timepoints. For the chemical treatments, the above-mentioned differentiation and maintenance cocktails were supplemented with the recombinant IGFBP2 protein at 2nM (R&D, #674-B2-025), recombinant IGF-I protein at 10nM (Sigma, #I3769), recombinant IGF-II protein at 10nM (R&D, #292-G2-050) and Echistatin 100 nM (R&D, #3202). Chemicals were added to both induction and maintenance cocktails except for Echistatin which was added to the induction cocktail only and withdrawn 48h after induction since inhibiting the integrin receptor resulted in cell detachment when Echistatin was kept in culture for longer periods than 48h. In the Echistatin mixed with IGFBP2 condition, only IGFBP2 was kept after 48h. IGFBP2, IGF-I and IGF-II were first titrated at the concentrations shown in **Figure S7 D-E**.

### Cell proliferation assay

Sorted cells were split into four and seeded in 4 different wells of a 12well plate and allowed to attach and start to proliferate for 7 to 10 days. One well of each cell population was trypsinized after this period. Cells were resuspended in 1 ml of medium, counted twice using a hematocytometer, and the mean count was used as the baseline number of cells from which cell increase was calculated. The same counting was performed on the remaining wells every two days. The expansion medium was refreshed every two days.

### Mixing and transwell experiments

For the mixing experiments, unexpanded Lin– SVF cells were isolated with MACS using Miltenyi LD columns (Miltenyi, #130-042-901) on manual mono-MACS separators after staining with magnetic anti-human CD45 and CD31 microbeads (Miltenyi, #130-045-801 and #130-091-935) according to the manufacturer’s protocol. MACS-isolated Lin-cells from SC, OM, and PR samples were counted in duplicates and mixed at high density (65k cells /cm^2^) in 11 ratios from 0 to 100%. After 24h, the cells were induced to differentiate following the adipogenic differentiation protocol. For the transwell experiments, we used 96well plate format transwell inserts with 0.4 μm (Corning #CLS3391) pores to allow protein and small molecule diffusion through the membrane, but not cell migration. 96well transwell-receiving plates (Corning #3382) were first coated with type I collagen (Corning #354249) 1:500 in DPBS before use to facilitate cell adhesion. Sorted donor OM subpopulations and expanded receiver SC and PR SVF-adherent cells were plated and expanded separately onto the top transwell insert and the bottom receiving plate, respectively. When confluent, the transwell insert was put in contact with the receiver plate, and all cells were induced to differentiate following the listed differentiation protocol.

### Enzyme-linked immunosorbent assay (ELISA)

For the supernatant measure, cells were expanded for two passages and seeded into a 6well plate. Once confluent, the expansion medium was aspirated, and wells were washed twice with PBS to ensure residual serum, dead cell and protein removal. 2ml of OPTI-Pro serum-free medium (Thermo, #12309050) was added to each well and incubated with the cells at 37°C for 48h. After incubation, SFM medium was harvested, spun for 10 min at 4°C max speed to clear potential cell debris. Cleared supernatant was aliquoted and stored at – 80°C until further usage. For the whole AT IGFBP2 secretion assays, three times 200-400 mg of OM AT were put in 500μl of DPBS (Gibco #14190169) and incubated at 37°C for 24, 48 and 72 hours. After incubation, DPBS was harvested, spun for 10 min at 4°C max speed to clear potential cell debris and stored at -80°C until further usage. The Anti-human IGFBP2 ELISA kit (Sigma, #RAB0233-1KT) was used to quantify IGFBP2 protein in the supernatants according to the manufacturer’s recommendations. Before loading samples on the ELISA membranes, the total protein concentration was quantified using the Qubit™ Protein Broad Range assay kit (Thermo, #A50669) and 300 ng of total protein was added per reaction. Incubation of samples with primary antibodies was performed O/N at 4°C. At the end of the assay, absorbance was read at 450 nm using a SPARK® Microplate reader.

### Immunohistochemistry

Human AT biopsies were washed twice in PBS to remove excess blood and divided in 50 to 100 mg for fixation in 4% PFA (paraformaldehyde, electron microscopy grade (VWR #100504-858)) for 2 hours at 4°C with gentle shaking. Next, the tissue was washed with PBS and incubated with 30% sucrose O/N at 4°C with gentle shaking. Cryoblocks were prepared using Cryomatrix (Thermo Fisher Scientific #6769006), and 25-μm sections were generated using a Leica CM3050S cryostat at −30°C. The tissue was air-dried for 30 minutes at -20°C in the cryostat itself, then 1h at RT. Slides were additionally fixed 10 min in 4% PFA at RT, washed two times 5 minutes with PBS, permeabilized at RT with 0.25% TritonX100 (Sigma #T9284) for 10 minutes, washed twice with PBS again and antigen blocking was performed at RT for 30 minutes with 1% BSA in PBS. Primary antibodies (anti-TM4SF1, anti-MSLN, anti-PLIN1) in 1% BSA were applied O/N at 4°C with gentle shaking following the titrations indicated in **Supp. Table 5**. The following day, after two PBS washes, and quick 1% BSA dip, the secondary antibody (anti-rabbit AF-647) in 1% BSA was applied for 40 minutes at RT following the titrations in **Supp. Table 5**. Nuclei were stained with 1μg/ml DAPI (Sigma #D9564) for 10 minutes and washed twice in PBS prior to mounting with Fluoromount G (Southern Biotech #0100-01). The slides were then imaged with a Leica SP8 Inverted confocal microscope (objectives: HC PL Fluotar 10x/0.30 air, HC PL APO 20x/0.75 air, HC PL APO 40x/1.25 glyc, HC PL APO 63x/1.40 oil). The results presented in **Figures 5K-L** were replicated in at least three independent experiments. We note that we also verified that the signal we detected is not the result of autofluorescence of the AT or from unspecific binding of secondary antibodies (**Figure S5D**).

### Imaging and quantification of *in vitro* adipogenesis

On the 14^th^ day of differentiation, cells were either fixed with 4% PFA (EMS, #15710) and stained at a later timepoint or live-stained with fluorescence dyes: Bodipy 10 μg/ml (boron-dipyrromethene, Invitrogen #D3922) for lipids and Hoechst 1 μg/ml (Sigma, #B2883) for nuclei. Cells were incubated with the dyes in PBS, for 30 min in the dark, washed twice with PBS, and imaged. If the imaging was performed on live cells, we used FluoroBrite DMEM (Gibco # A1896701) supplemented with 10% FBS as acquisition medium. Given substantial variation in the extent of lipid accumulation by the tested cell fractions (within the same well but also across technical replicates), the imaging was optimized to cover the largest surface possible of the 96 well. Moreover, a z-stack acquisition in a spinning-disc mode and Z-projection were performed in order to capture the extent of *in vitro* adipogenesis with the highest possible accuracy. Specifically, the automated platform Operetta (Perkin Elmer) was used for imaging. First, 3–6 z-stacks were acquired for every field of view in a confocal mode of the microscope in order to produce high-quality images for downstream z-projection and accurate thresholding. Next, 25 images per well were acquired using a Plan Neofluar 10× Air, NA 0.35 objective for the transwell-receiving plates or 20x air objective NA 0.8 for normal 96w plates (Falcon, #353219), with no overlap for further tiling and with the aim of covering the majority of the well for an accurate representation of lipid accumulation (see Methods in ^17^). The lasers were set in time exposure and power to assure that in both the Hoechst and the Bodipy channels, the pixel intensity was between 500 and 4000, and in all cases at least two times higher than the surrounding background. The images, supported by Harmony software, were exported as TIFF files. They were subsequently tiled, and Z-projected with the maximum intensity method. To accurately estimate and represent differences in adipocyte differentiation, a quantification algorithm for image treatment was developed in collaboration with the EPFL BIOP imaging facility. In brief, image analysis was performed in ImageJ/Fiji, lipid droplets (yellow) and nuclei (blue) images were filtered using a Gaussian blur (sigma equal to 2 and 3, respectively) before automatic thresholding. The automatic thresholding algorithm selections were chosen based on visual inspection of output images. The area corresponding to the thresholded lipid signal was then divided by the area corresponding to the thresholded nuclei area and used to calculate the Adiposcore (totalLipidArea/totalNucleiArea). In the figures, representative blown-up cropped images of each sample are shown. To reduce technical variation across the biological replicates (different donors), adiposcores were normalized to the average adiposcore of the indicated control when we compared conditions within highly differentiating lines like SC and PR. Adiposcores were compared without normalization when we wanted to directly compare adiposcores across depots (**Figures 1C**, **S1B**) or among poorly-differentiating samples like OM when the absolute values of adiposcores were < 0.01 (**Figures 4F, 5E, 7J, 7N-P**).

### siRNA-mediated knockdown

To achieve knockdown of IGFBP2, direct transfection was performed on OM SVF Lin–/TM4SF1+/MSLN– cells using the IGFBP2 IDT, TriFECTA DsiRNAs kit using 3 pooled siRNAs: hs.Ri.IGFBP2.13.1, hs.Ri.IGFBP2.13.2, hs.Ri.IGFBP2.13.3. In brief, after sorting, cells were expanded for one or two rounds, then harvested and plated at mid-low density (45k cells/cm2) and allowed to adhere. The following day, transfection mix was prepared as Opti-MEM medium (Invitrogen #31985062), 1.5% Lipofectamine RNAiMAX (Invitrogen #13778150) and 20 nM of the pooled siRNAs. In the transfection mix, lipofectamine-siRNA transfection particles were allowed to form for 15 min at RT with gentle shaking. After incubation, the transfection mix was diluted 10 times (to a final concentration of siRNA of 2 nM) in MEMalpha GlutaMax medium (Gibco #32561037) supplemented with 2.5% human platelet lysate (Sigma #SCM152), w/o antibiotics and exchanged to the plated cell medium. After 48h, medium was changed to differentiation medium (for the transwell assay), with serum free medium (for ELISA validation) or directly taken in TRIzol (for qPCR validation).

### RNA isolation and qPCR

Expanded OM and SC SVF-adherent, OM SVF Lin–/TM4SF1–/MSLN–, OM SVF Lin–/TM4SF1+/MSLN– cells as well as cells subjected to siRNA-mediated knockdowns 48h post-transfection were collected into TRIzol (Sigma, #T3934). The direct-zol RNA kit (Zymo Research #R2062) was used to extract RNA, followed by reverse transcription using the SuperScript II VILO cDNA Synthesis Kit (Invitrogen # 11754050). Expression levels of mRNA were assessed by real-time PCR using the PowerUp SYBR Green Master Mix (Thermo Fisher Scientific #A25743). mRNA expression was normalized to the Hprt1 gene. Primer sequences used: *IGFBP2* – Fw CGAGGGCACTTGTGAGAAGCG, Rv TGTTCATGGTGCTGTCCACGTG; *HPRT* – Fw CAGCCCTGGCGTCGTGATTA, Rv GTGATGGCCTCCCATCTCCTT.

### Statistical methods

The experiments were not randomized, and the investigators were not blinded in experiments. The paired Student’s *t*-test was used to determine statistical differences between two groups, with the null hypothesis being that the two groups are equal. Multiple comparisons were corrected using false discovery rate (FDR) correction. When specified, one-way ANOVA or RELM test followed by Tukey honest significant difference (HSD) post hoc correction was applied, the null hypothesis being defined so that the difference of means was zero. (Adjusted) \**p*-value < 0.05, \*\**p*-value < 0.01, \*\*\**p*-value < 0.001 were considered statistically significant. All boxplots display the mean as a dark band, the box shows the 25^th^ and 75^th^ percentiles, while the whiskers indicate the minimum and maximum data points in the considered dataset excluding outliers. All bar plots display the mean value and the standard deviation from the mean as error bar.

## Acknowledgment

The authors thank the EPFL and UNIL Core Facilities: FCCF (Flow Cytometry Core Facility EPFL, especially Miguel Garcia), FCF (Flow Cytometry Facility, especially Danny Labes), BIOP (BioImaging and Optics Platform, especially Olivier Buri, Romain Guiet and José Artacho), GECF (Gene Expression Core Facility, especially Bastien Mangeat and Elisa Cora), HCF (Histology Core Facility, especially Jessica Dessimoz). Open access funding provided by EPFL, a Leenaards Foundation MD-PhD Fellowship to R.F. (#531466) and a Leenaards Prize for Biomedical Translational Research to B.D. (Project #5154.5), a Personalized Health and Related Technologies (PHRT) PhD student fellowship (#2017/307) to P.R. and PHRT Grants (#2017/502 & #2019/719) to B.D., as well as a Swiss National Science Foundation Project grant (#310030_182655) to B.D.

## Authors contributions

R.F., P.R. and B.D. designed the study and wrote the manuscript. R.F. conducted all experimental procedures and analyzed acquired images, flow cytometric measures, qPCRs, ELISAs and immunohistochemistry. P.R. conducted all analyses related to transcriptomics both at the single-cell and bulk levels. J.R. and M.Z. assisted with sample processing, cell culture and preparation of sequencing libraries. J.R. performed histological assays. J.P. provided assistance with flow cytometry-related procedures. D.A. and V.G. assisted with bulk transcriptomic-associated procedures and data processing. V.G. and W.S. assisted with the transcriptomic analyses. L.F., S.M., T. Z., N.P., M.S. and M.M. provided access to human samples. V.G., W.S. and C.C. provided extensive comments to the manuscript.

## Corresponding author

Bart Deplancke

