## Supplementary Figures for "A human omentum-specific mesothelial-like stromal population inhibits adipogenesis through IGFBP2 secretion": All Supp. Figures.pdf

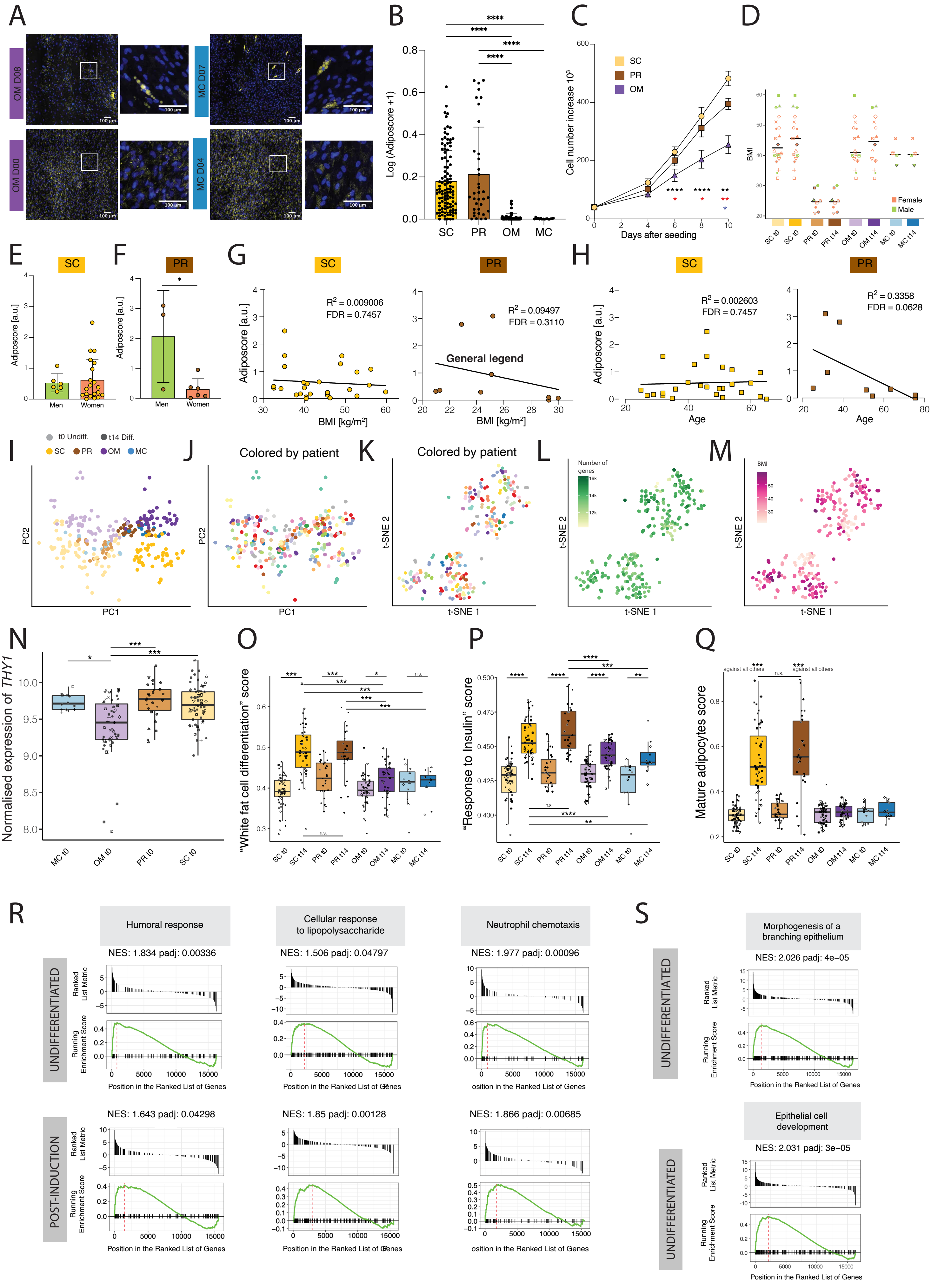

**Figure S1** | See next page for caption

**Figure S1. Comparisons of primary SVF-adherent cells across depots.**

- **(A)** Representative confocal images of intraperitoneal omentum (OM)- and mesocolic (MC)-derived SVF-adherent cells after 14 days of differentiation; Top: lines that form very few mature lipid droplets, Bottom: lines that form small lipid droplets that are barely distinguishable from background; Yellow - Bodipy staining for lipids, blue - Hoechst staining for DNA.
- **(B)** Cross-anatomical comparison of the adiposcores of differentiated SVF-adherent lines from **Figure 1D** and sequenced in **Figure 1E**. On the y axes, the log(adiposcore+1) is plotted. Subcutaneous (SC): n=104, 4 independent wells, 26 cell lines, 20 donors; Perirenal (PR): n=36, 4 independent wells, 9 cell lines, 8 donors; OM: n=88, 4 independent wells, 22 cell lines, 18 donors; MC: n=16, 4 independent wells, 4 cell lines, 4 donors.
- **(C)** Relative cell numbers over time of culture for SC, OM, and PR SVF-adherent cells; n=12, 4 donors per depot; Black stars compare SC *versus* OM, Red: PR *versus* OM, Blue: SC *versus* PR.
- **(D)** Plot showing the number of donors and distribution of their BMI and sex included in the BRB-seq[20] analysis across the different depots and time points, shown in **Figure 1E**.
- **(E)** Bar plot showing the distribution of the adiposcore of SC SVF-adherent lines between men and women.
- **(F)** Bar plot showing the distribution of the adiposcore of PR SVF-adherent lines between men and women.
- **(G)** Scatter plot showing the correlation between the adiposcore of highly adipogenic SVF-adherent lines (SC - **left** and PR - **right**) and the BMI of respective donors. The line represents a linear regression analysis.
- **(H)** Scatter plot showing the correlation between the adiposcore of highly adipogenic SVF-adherent lines (SC - **left** and PR - **right**) and the age of respective donors. The line represents a linear regression analysis.
- **(I)** PCA based on the BRB-seq[20] data of SVF-adherent cells from the indicated adipose depots at different time points (t0 - undifferentiated, t14 - 14 days post-adipogenic induction).
- **(J)** PCA, as described in **I**, colored by donors.
- **(K)** t-SNE map shown in **Figure 1E** computed on the 10 first PCs of the PCA displayed in **I**, colored by donors.
- **(L)** t-SNE map shown in **Figure 1E** colored by the number of detected genes.
- **(M)** t-SNE map shown in **Figure 1E** colored by the BMI of the donors.
- **(N)** Boxplot displaying the expression distribution of *THY1*, a known mesenchymal cell marker, across samples from the indicated depots at t0.
- **(O)** Box plots displaying the “white fat cell differentiation score” based on the scaled expression of the corresponding GO term (GO:0050872) of the data shown in **Figure 1E**.
- **(P)** Box plots displaying the “response to insulin score” based on the scaled expression of the corresponding GO term (GO:0032868) of the data shown in **Figure 1E**.
- **(Q)** Box plots displaying the “mature adipocyte score” based on the scaled expression of the following markers: *FABP4, PPARG, ADIPOQ, LIPE, LPL, PLIN1, PLIN2, PLIN4, CEBPA, CEBPB, CIDEC,* and *CIDEA* for the indicated depots and time points of the data shown in **Figure 1E**.
- **(R)** GSEA plot of selected inflammatory response GO terms (“humoral response” GO:0006959, “cellular response to lipopolysaccharides” GO:0071222, “neutrophil chemotaxis” GO: 0030593), based on the differential expression analysis of SVF-adherent cells derived from OM adipose depots *versus* those from other depots (SC, PR, MC) at t0 (i.e., undifferentiated state) or t14 (i.e., post-adipogenic induction).
- **(S)** GSEA plot of the GO terms “epithelial cell development” (GO:0060429) and “morphogenesis of a branching epithelium” (GO:0048754), based on the differential expression analysis of

SVF-adherent cells derived from OM adipose depots *versus* those from other depots (SC, PR, MC) at t0 (i.e., undifferentiated state).

*p 0.05, **p 0.01, ***p 0.001, ****p 0.0001, One-Way ANOVA and Tukey HSD *post hoc* test (**B**, **E**, **F**), and RELM analysis and Tukey HSD *post hoc* test (**C**), linear regression analysis with its relative goodness of fit, and the FDR-adjusted *p*-values of the Pearson correlations (**G**, **H**), unpaired two-sided *t*-test (**N**-**Q**).

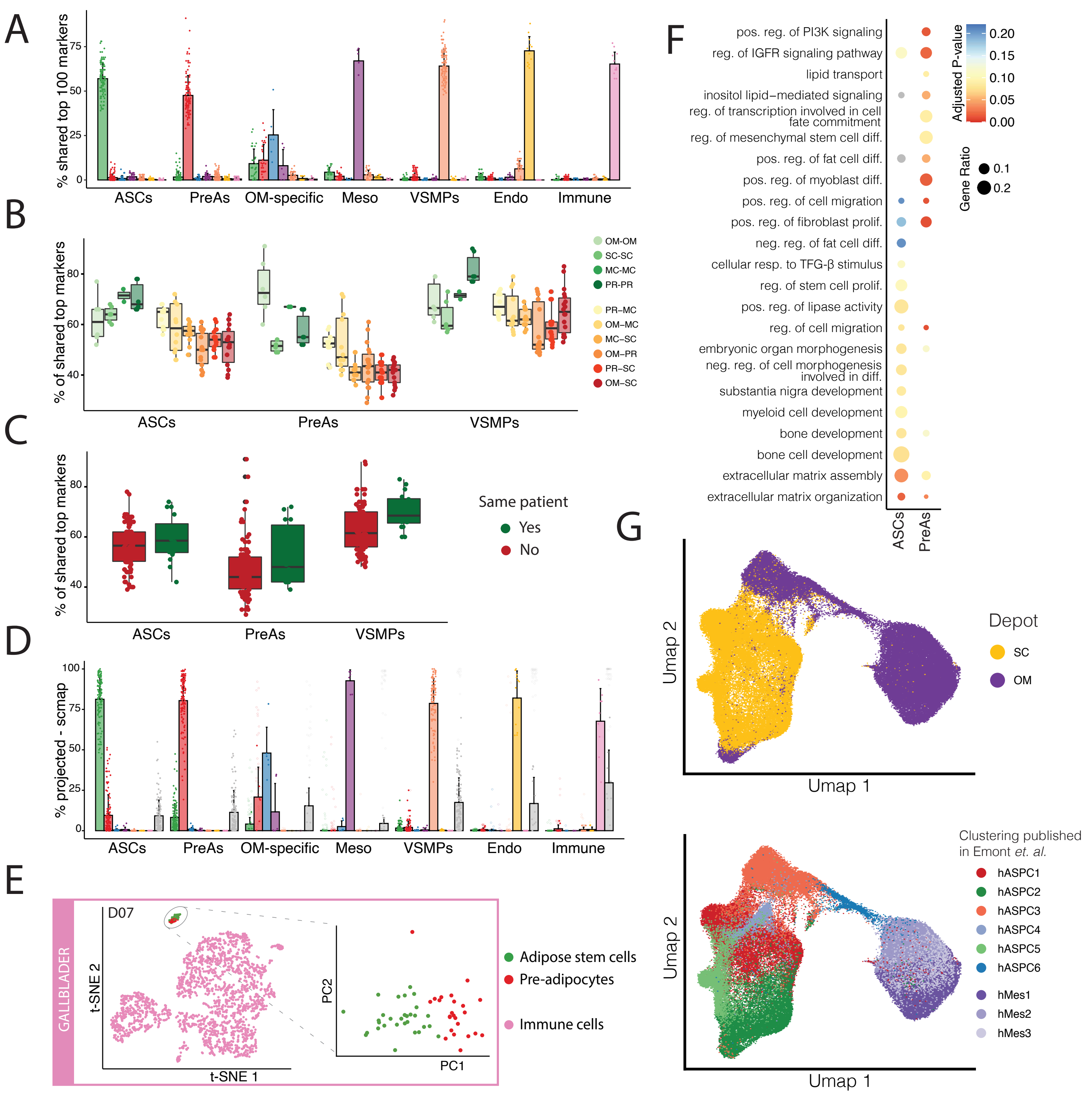


**Figure S2 |** See next page for caption

**Figure S2. Human adipose-derived stromal cells feature an anatomic footprint in their transcriptome.**

- **(A)** The percentage of shared top 100 differentially expressed genes between each subpopulation and sample shown in **Figure 1A**; each point represents the number of shared markers between the indicated subpopulations (x-axis) of an individual dataset X and a subpopulation (coloring) in another individual dataset Y.
- **(B)** The percentage of shared top 100 differentially expressed genes as shown in panel **A** but limited to the comparison of the same populations (ASCs with ASCs, PreAs with PreAs, and VSMPs with VSMPs) across depots and donors, split by the type of depot of the comparison pairs; comparisons between samples originating from the same depots are highlighted in shades of green, and comparison of samples originating from different depots are in shades from yellow to red.
- **(C)** The percentage of shared top 100 differentially expressed genes as shown in panel **A** but limited to the comparison of the same populations (ASCs with ASCs, PreAs with PreAs, and VSMPs with VSMPs) across depots and donors, stratified according to whether the pairs of compared samples are originating from the same donor (green) or not (red).
- **(D)** The percentage of cells of a subpopulation projected onto each subpopulation and sample based on scmap[29] results (see **Methods**). Each point represents the percentage of a subpopulation (x-axis) of an individual dataset X projected onto a subpopulation (coloring) of another dataset Y. Projections of subpopulations (x-axis) non-existing in the reference data are highlighted as shaded circles.
- **(E)** **left** - t-SNE cell map of a scRNA-seq dataset of SVF Lin– cells isolated from gallbladder-associated adipose tissue from one donor colored by populations. **right** - PCA of the highlighted hASPCs of the left panel visualizing adipose stem cells (ASCs; green) and pre-adipocytes (PreAs; red), a total of 54 cells.
- **(F)** Dot plot showing enriched biological process GO terms based on differentially expressed genes of ASCs or PreAs of the integrated scRNA-seq data shown in **Figure 2C**.
- **(G)** UMAP of hASPCs and human mesothelial cells from scRNA-seq data provided by Emont et al.[8] colored by the depot of origin (**top**) or colored by the clustering published in the latter study (**bottom**).

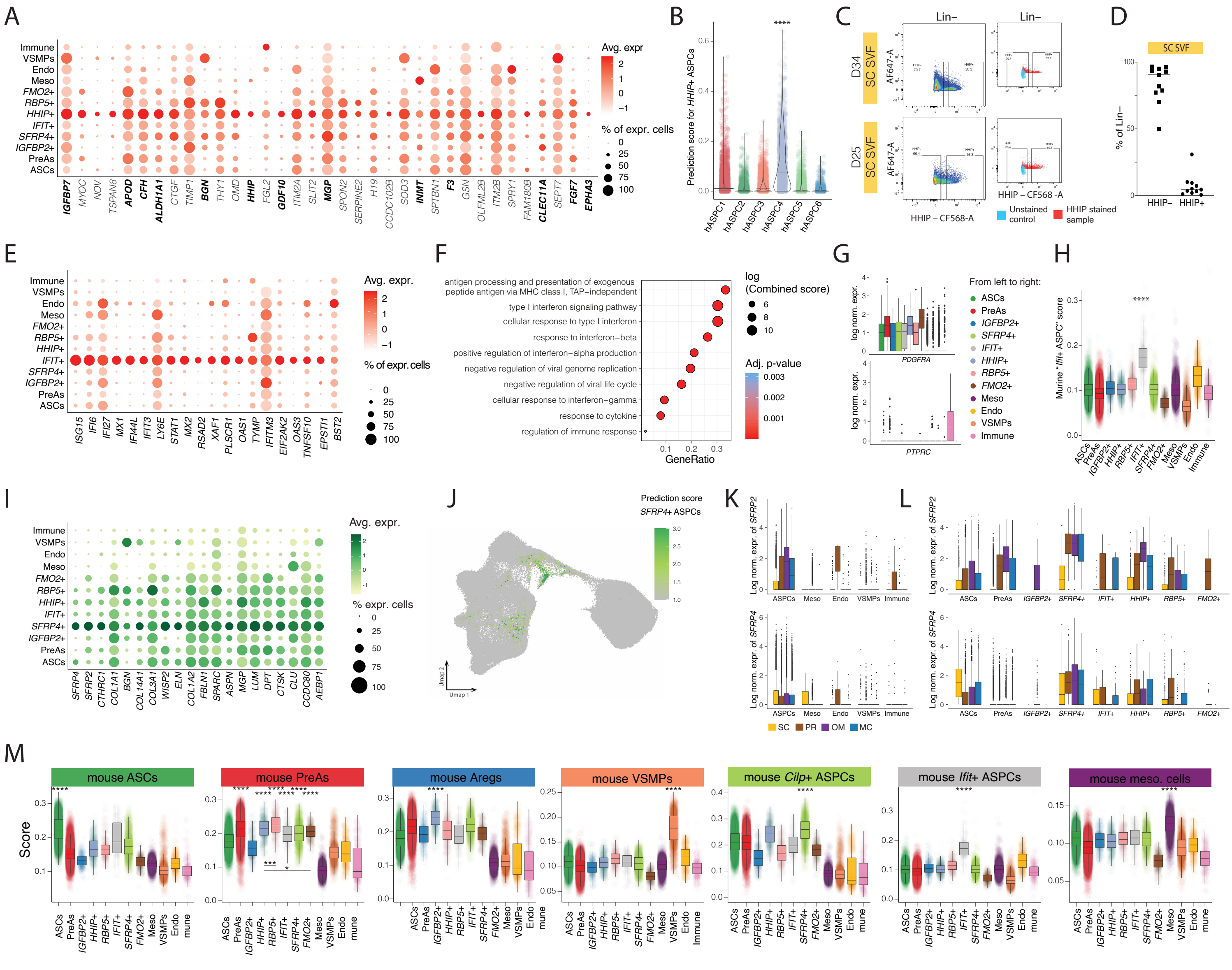

**Figure S3 |** See next page for caption

**Figure S3. Molecular identity of minor human stromal cells.**

- **(A)** Dot plot displaying the average expression and percentage of expressing cells of the top 20 markers of *HHIP*+ hASPCs across the clusters shown in **Figure 2C**. The orthologs of the murine top Areg markers (as defined in Zachara et al.[17]) are highlighted in bold.
- **(B)** Violin plot showing the distribution of the prediction score of the *HHIP*+ hASPC population when transferred onto the scRNA-seq atlas of hASPCs of Emont et al.[8], where they identified hASPC4 as being transcriptionally similar to murine Aregs[12,17].
- **(C)** Representative flow cytometry-based gating of HHIP+ events on subcutaneous (SC) adipose depot-derived SVF Lin– cells.
- **(D)** Flow cytometry-based quantification of Lin–/HHIP+ events in SC SVF Lin–, n=11.
- **(E)** Dot plot showing the average expression and percentage of expressing cells of the top 20 markers of *IFIT*+ hASPCs across the clusters shown in **Figure 2C**.
- **(F)** Dot plot showing representative GO terms that are enriched based on the differentially expressed genes of *IFIT*+ hASPCs.
- **(G)** Box plots showing the distribution of the log normalized expression of *PDGFRA* and *PTPRC* (*CD45*) across the cluster shown in **Figure 2C**.
- **(H)** Box plot showing the distribution of the murine "*Ifit*+ ASPC" scores across the detected, distinct human SVF cell populations. The scores were based on the human orthologs of the murine top markers of the *Ifit*+ ASPCs based on the integration of scRNA-seq datasets of subcutaneous and visceral murine adipose tissues described in Ferrero et al.[11].
- **(I)** Dot plot displaying the average expression and percentage of expressing cells of the top markers of *SFRP4*+ hASPCs across the clusters shown in **Figure 2C**.
- **(J)** UMAP of hASPCs and human mesothelial cells from scRNA-seq data provided by Emont et al.[8] colored by the prediction score of *SFRP4*+ cells when transferring our cell cluster annotation.
- **(K)** Box plots showing the distribution of *SFRP2* (**top**) and *SFRP4* (**bottom**) expression in the different cell types from the integrated scRNA-seq data, colored by depot of origin.
- **(L)** Box plots showing the distribution of *SFRP2* (**top**) and *SFRP4* (**bottom**) expression in the different hASPC subpopulations from the integrated scRNA-seq data, colored by depots of origin.
- **(M)** Box plots showing for each mouse cell population defined in Ferrero et al.[11] the score of orthologous human markers in each human cell population identified in **Figure 2C**.

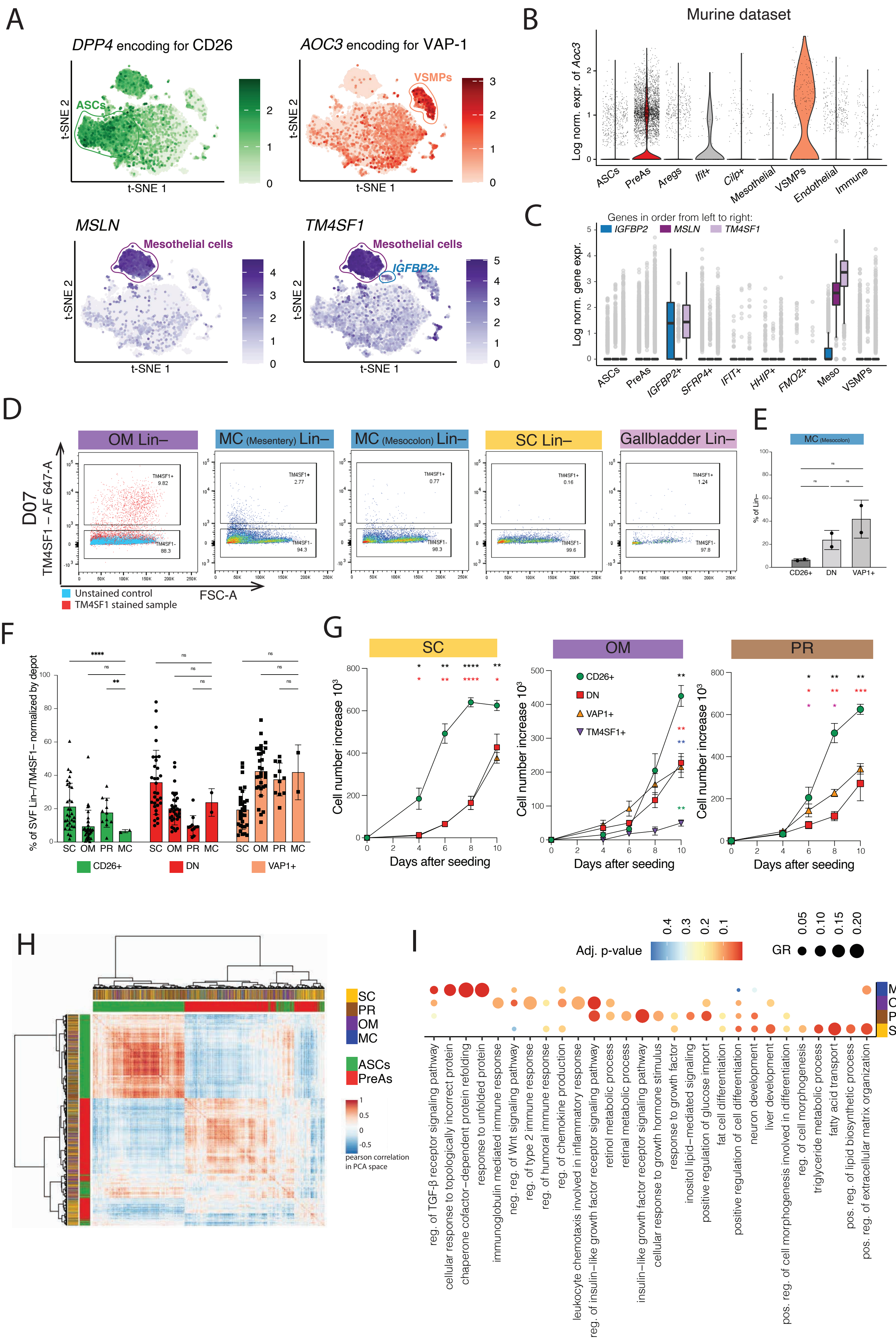


**Figure S4 |** See next page for caption

**Figure S4. The selection of scRNA-seq-inferred surface markers enables the enrichment of the main SVF cell populations across all analyzed adipose depots.**

- **(A)** t-SNE cell map of integrated scRNA-seq datasets described in **Figure 2C** colored by the expression of genes corresponding to the surface markers that were used to isolate each subpopulation experimentally.
- **(B)** Violin plot showing the distribution of the log normalized expression of *Aoc3* based on the scRNA-seq integration of murine datasets (see **Methods**).
- **(C)** Box plot showing the distribution of the log normalized scRNA-seq-based expression for *IGFBP2*, *MSLN*, and *TM4SF1* (color) across the different human cell populations (x-axis).
- **(D)** Flow cytometry profiles of Lin–/TM4SF1+ populations after TM4SF1 staining from five different adipose depot-derived SVF cells from the same donor.
- **(E)** Flow cytometry-based analysis of the abundance of each indicated cell population gated from the Lin– fraction of MC SVF cells. Bar plots indicate mean, error bars standard deviation; n=2 donors.
- **(F)** Bar plot to compare flow cytometry-based abundances of the indicated cell populations across subcutaneous (SC), perirenal (PR), omentum (OM), and mesocolic (MC) adipose depots within the SVF Lin– fraction. The three populations accumulate to 100% by depot. For OM, each cell population is also TM4SF1– to deplete for OM-specific populations; SC n=37, OM n=35, PR n=17; MC n=2 donors.
- **(G)** Relative cell number increase over time of culture for CD26+, DN, and VAP1+ cells in each depot. For OM, the three cell populations were gated from the Lin–/TM4SF1– population and the proliferation of Lin–/TM4SF1+ OM-specific cells was also recorded; n=12, 3 donors, 3- 4 populations per depot.
- **(H)** Heatmap of the correlation between 2000 randomly selected ASCs and PreAs based on the first 30 principal components of the PCA space of integrated scRNA-seq data shown in **Figure 2C**; a similar number of cells was selected for each depot and population.
- **(I)** Dot plot of enriched, representative GO terms based on the differentially expressed genes specific to the indicated depot, as explained in **Figure 4H**.

**p* 0.05, ***p* 0.01, ****p* 0.001, *****p* 0.0001, One-Way ANOVA and Tukey HSD *post hoc* test (**E**, **F**) and RELM analysis and Tukey HSD *post hoc* test (**G**). Black compares CD26+ *versus* DN, Red CD26+ *versus* VAP1+, Blue DN *versus* TM4SF1+, Green VAP1+ *versus* TM4SF1+, Pink DN *versus* VAP1+.

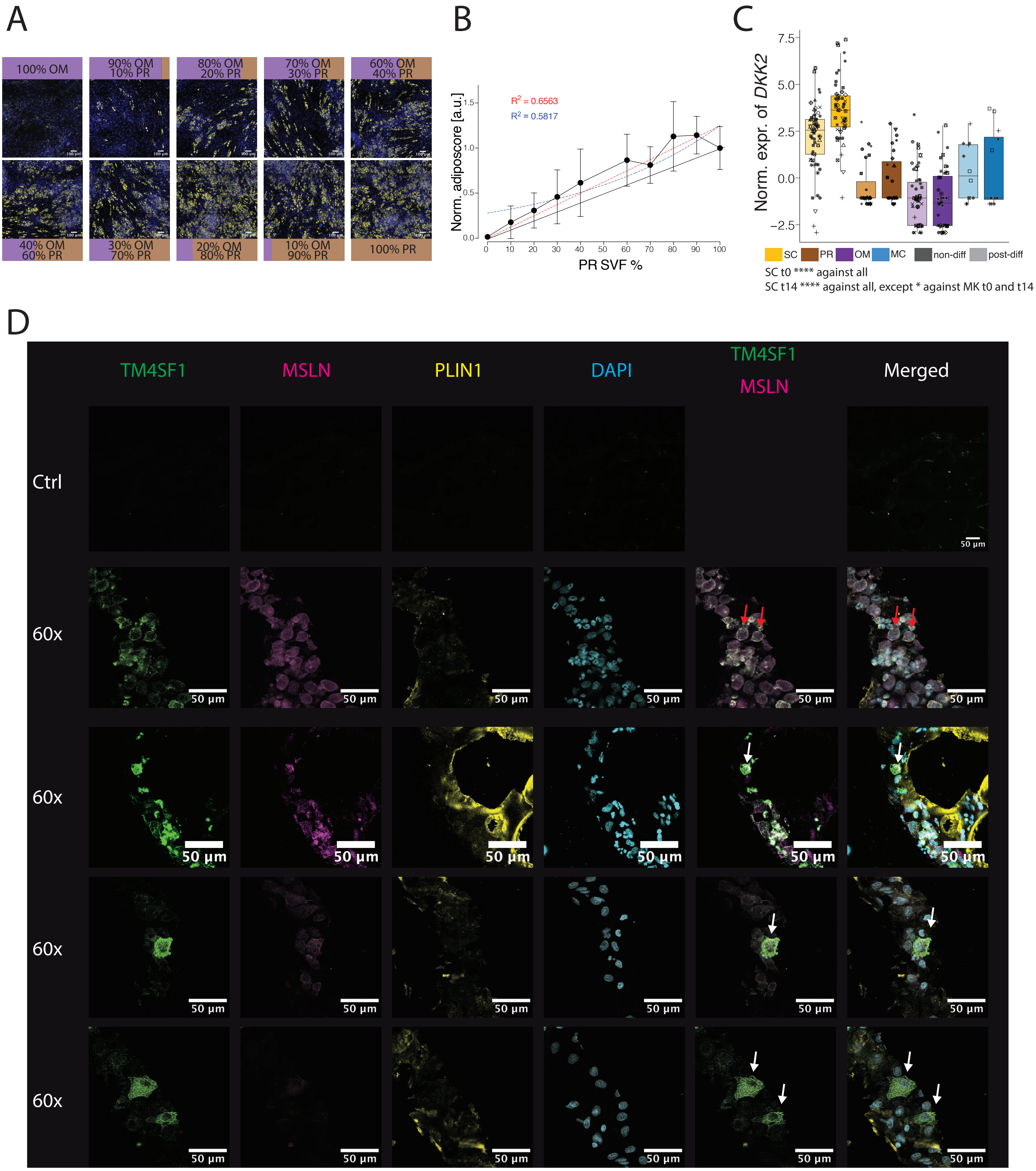


**Figure S5 |** See next page for caption

**Figure S5. OM inhibitory cells act in a depot-specific way.**

**(A)** Representative fluorescence microscopy images of SVF Lin– cells in mixing experiments after 14 days of adipogenic differentiation, where SVF Lin– cells from omentum (OM) and perirenal (PR) fat of donor 68 were mixed directly after cell isolation at the indicated proportions. Yellow - Bodipy staining for lipids, blue - Hoechst staining for DNA, scale bar=100 µm.

**(B)** Adiposcore of the mixed OM and PR SVF Lin– cell populations, as presented in **A**. Values across biological replicates are normalized to the average adiposcore of the reference 100% PR Lin– condition. The relative proportion (0-100%) of PR SVF Lin– cells in each well is plotted on the x-axis. Error bars represent standard deviation from the average, linear and exponential regression with corresponding $R^2$ coefficients are shown in red and blue, respectively. The black line represents the expected increase of adipogenesis for a linear dilution between 0 and 100% of PR SVF Lin– cells; n=16, 4 biological replicates, 4 independent wells for each.

**(C)** Box plot showing the distribution of batch normalized expression of DKK2 of BRB-seq[20] data of SVF-isolated cells from the indicated depots and treatment conditions, n=12-61, 4-20 biological replicates, 1-4 independent wells for each.

**(D)** Confocal microscopy fluorescent images after TM4SF1 (green), Perilipin (PLIN1) (yellow) and MSLN (pink) immunohistochemistry staining of whole OM AT cryocuts. The top row is the unstained control. DAPI staining for nuclei is colored in cyan. The experiment was repeated three times. The white arrows point to TM4SF1+ cells, the red arrows point to TM4SF1+/MSLN+ cells.

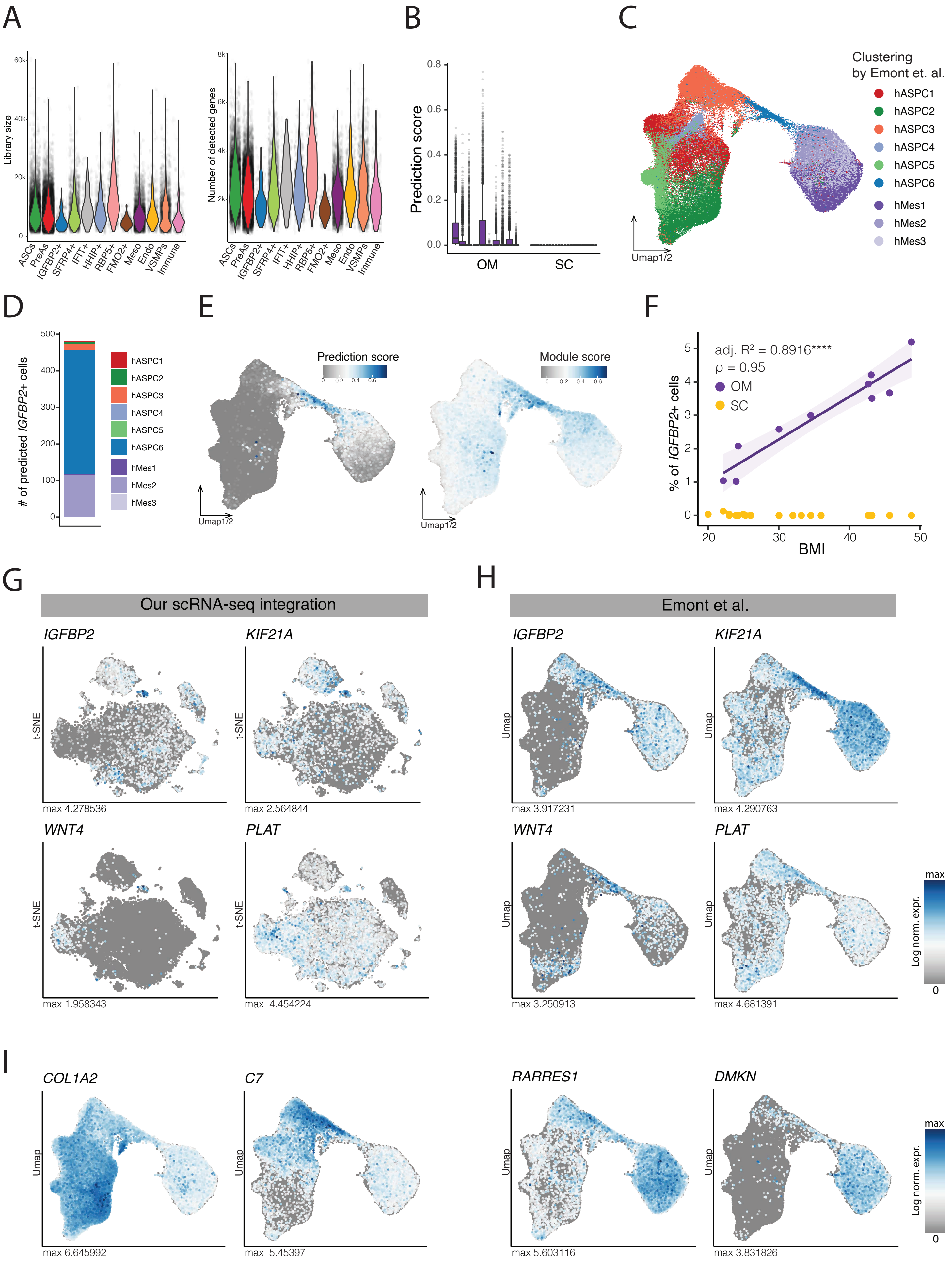


**Figure S6 |** See next page for caption

**Figure S6. Identification of omental IGFBP2+ cells using computational methods and independently published datasets.**

- **(A)** Violin plots showing the distribution of library size (**left**) or the number of detected genes (**right**) across the different clusters shown in **Figure 2C**.
- **(B)** Box plot showing the distribution of the prediction score of *IGFBP2*+ cells when transferring our cell cluster annotation on the data published by Emont et al.[8] for the indicated adipose depots (x-axis); SC - Subcutaneous, OM - Omentum.
- **(C)** UMAP computed on the integrated data of hASPCs and human mesothelial cells reported by Emont et al.[8] colored by the clustering provided in the same study.
- **(D)** Bar plot displaying the number of cells predicted as *IGFBP2*+ cells into each of the clusters of mesothelial cells and ASPCs originally reported by Emont et al.[8], shown in **C**.
- **(E)** UMAP described in **C** colored by the prediction score of *IGFBP2*+ cells when transferring our cell cluster annotation on the data reported by Emont et al.[8] (**left**), or colored by the score based on the top *IGFBP2*+ cell markers (**right**).
- **(F)** Correlation between every donor's BMI and the percentage of hASPC6 cells (*IGFBP*2+-like) for each donor based on the scRNA-seq dataset provided by Emont et al.[8]; the percentage was calculated for each donor as the fraction of mesothelial cells and ASPCs combined.
- **(G)** t-SNE cell map of our integrated scRNA-seq data colored by the log-normalized expression of the indicated *IGFBP2*+ cell markers.
- **(H)** UMAP described in **C** colored by the log-normalized expression of some *IGFBP2*+ cell markers as in **G**.
- **(I)** UMAP described in **C** colored by the log-normalized expression of the indicated markers shared by predicted *IGFBP2*+ cells and ASPCs (two **left** plots) or Mesothelial cells (two **right** plots).

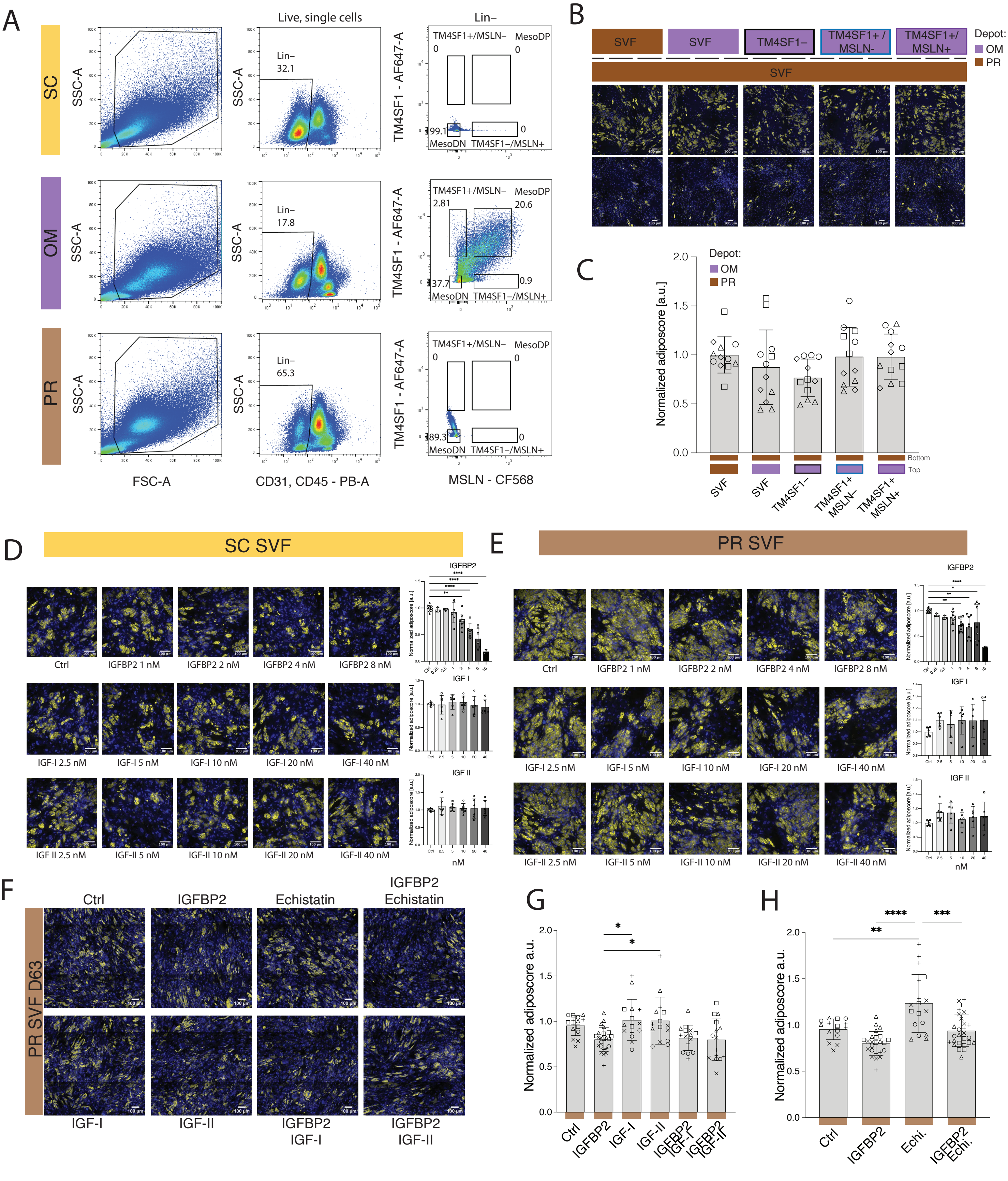


**Figure S7 |** See next page for caption

**Figure S7. The *IGFBP2*+ cell-mediated inhibition is less effective upon PR-derived lines.**

- **(A)** Representative flow cytometry plots of SC (Subcutaneous), OM (Omentum), and PR (Perirenal) SVF/Lin– from donor 53 stained with TM4SF1 and MSLN and gating strategy to enrich for ASPCs (Lin–/TM4SF1–/MSLN–), *IGFBP2*+ cells (Lin–/TM4SF1+/MSLN–) or mesothelial cells (Lin–/TM4SF1+/MSLN+) exclusively in OM SVF. Similar profiles were obtained from at least three donors.
- **(B)** Representative fluorescence microscopy images of "receiver" PR SVF-adherent cells, at the bottom of the transwell set-up, after adipogenic differentiation when co-cultured with the indicated SVF fractions at the top: paired PR SVF-adherent cells, OM SVF-adherent cells, OM SVF/Lin–/TM4SF1– (OM ASPCs), OM SVF/Lin–/TM4SF1+/MSLN– (IGFBP2-secreting cells), or OM SVF/Lin–/TM4SF1+/MSLN+ cells (mesothelial cells). First row: PR and OM cells from D54, Second row: SC and OM cells from D65.
- **(C)** Bar plot showing the adiposcore quantification of bottom cells in **B**. Values are normalized to the average adiposcore of the reference top PR SVF-adherent condition; n=12, 4 donors, 3 independent wells.
- **(D)** Representative fluorescent microscopy images and adiposcore quantification of SC SVF-adherent cells treated with the indicated compounds and concentrations. Scale bars=100µm. n=3-10, 3 donors, 2-6 independent wells.
- **(E)** Representative fluorescent microscopy images and adiposcore quantification of PR SVF-adherent cells treated with the indicated compounds and concentrations. Scale bars=100µm. n=3-12, 4 donors, 2-6 independent wells.
- **(F)** Representative fluorescence microscopy images of PR SVF cells after adipogenic differentiation when treated with the indicated interfering compounds. IGFBP2 1nM, IGF-I 10nM, IGF-II 10nM, Echistatin 100nM; Scale bars=100µm.
- **(G)** Barplot showing the adiposcore quantification of cells in **F** focusing on the IGF-dependent signaling pathway of IGFBP2. The adiposcores are normalized to the non-treated cells (Ctrl); n=12, 4 donors, 3 independent wells.
- **(H)** Barplot showing the adiposcore quantification of cells in **F** focusing on the IGF-independent signaling pathway of IGFBP2. The adiposcores are normalized to the non-treated cells (Ctrl). n=12, 4 donors, 3 independent wells.

SC - Yellow, OM - Purple, PR - Brown; For fluorescence images: Yellow - Bodipy staining for lipids, blue - Hoechst staining for DNA. **p* 0.05, ***p* 0.01, ****p* 0.001, *****p* 0.000,1. One-Way ANOVA and Tukey HSD *post hoc* test.
