## Supplementary Tables for "A human omentum-specific mesothelial-like stromal population inhibits adipogenesis through IGFBP2 secretion"

| Donor code | Fig1 & supp | Fig2 & supp | Fig3 & supp | Fig4 & supp | Fig 5 & supp | Height | Weight | BMI | Age | Gender | Type of surgery |
|---|---|---|---|---|---|---|---|---|---|---|---|
| D00 | CI, AS, BRB | scRNAseq | | | | 1.74 | 138 | 45.58 | 32 | Female | Gastric bypass |
| D01 | CI, AS, BRB | scRNAseq | | | | 1.77 | 110 | 35.11 | 46 | Female | Gastric bypass |
| D02 | CI, AS, BRB | | | | | 1.78 | 146.5 | 46.23 | 35 | Female | Gastric bypass |
| D03 | CI, AS, BRB | | | | | 1.64 | 108.6 | 40.37 | 51 | Female | Gastric bypass |
| D04 | CI, AS, BRB | | | | | 1.66 | 112 | 40.64 | 54 | Female | Gastric bypass |
| D05 | CI, AS, BRB | | FC | CI, AS | | 1.66 | 112 | 40.64 | 54 | Female | Gastric bypass |
| D06 | CI, AS, BRB | | FC | | | 1.71 | 111 | 37.96 | 27 | Female | Gastric bypass |
| D07 | CI, AS, BRB | scRNAseq | FC | | | 1.79 | 128 | 39.94 | 51 | Male | Gastric bypass |
| D08 | CI, AS, BRB | | | | | 1.59 | 129 | 51.02 | 25 | Female | Gastric bypass |
| D09 | CI, AS, BRB | | FC | | | 1.6 | 106 | 41.4 | 29 | Female | Gastric bypass |
| D10 | CI, AS, BRB | | FC | | | 1.53 | 102 | 43.57 | 40 | Female | Gastric bypass |
| D11 | CI, AS, BRB | | FC | | | 1.64 | 132 | 49.07 | 32 | Female | Gastric bypass |
| D12 | CI, AS, BRB | | FC | | | 1.76 | 106 | 34.22 | 36 | Male | Gastric bypass |
| D13 | CI, AS, BRB | | FC | | | 1.67 | 136 | 48.76 | 42 | Female | Gastric bypass |
| D14 | CI, AS, BRB | | FC | Mix AS | | 1.56 | 93 | 38.21 | 26 | Female | Gastric bypass |
| D15 | CI, AS, BRB | | FC | Mix AS | | 1.7 | 115 | 39.79 | 60 | Male | Gastric bypass |
| D16 | CI, AS, BRB | | FC | | | 1.79 | 180 | 56.17 | 54 | Male | Gastric bypass |
| D17 | CI, AS, BRB | | FC | | | 1.74 | 181 | 59.78 | 57 | Male | Gastric bypass |
| D18 | CI, AS, BRB | | | | | 1.72 | 96 | 32.44 | 50 | Female | Gastric bypass |
| D19 | CI, AS, BRB | | FC | | | 1.74 | 169 | 55.81 | 44 | Male | Gastric bypass |
| D20 | CI, AS, BRB | | FC | | | 1.68 | 149 | 52.79 | 63 | Female | Gastric bypass |
| D21 | CI, AS, BRB | | FC | | | 1.68 | 110 | 38.97 | 49 | Female | Gastric bypass |
| D22 | CI, AS, BRB | | FC | | | 1.89 | 90 | 25.19 | 31 | Male | Nephrectomy |
| D23 | CI, AS, BRB | | FC, Profiles | | | 1.6 | 75 | 29.29 | 75 | Female | Nephrectomy |
| D24 | CI, AS, BRB | scRNAseq | FC | | TW AS | 1.53 | 57 | 24.34 | 63 | Female | Nephrectomy |
| D25 | CI, AS, BRB | | FC | | | 1.62 | 55 | 20.95 | 65 | Female | Nephrectomy |
| D26 | | | FC | | | 1.61 | 102 | 39.35 | 60 | Female | Gastric bypass |
| D27 | CI, AS, BRB | | FC | | | 1.75 | 92 | 30.04 | 59 | Male | Nephrectomy |
| D28 | CI, AS, BRB | | | | | 1.61 | 65 | 25.07 | 45 | Female | Nephrectomy |
| D29 | CI, AS, BRB | | FC | | | 1.62 | 60 | 22.86 | 46 | Female | Nephrectomy |
| D30 | CI, AS, BRB | scRNAseq | FC, AS | | | 1.68 | 60 | 21.25 | 63 | Female | Nephrectomy |
| D31 | CI, AS, BRB | | | | | 1.76 | 114 | 36.8 | 38 | Male | Gastric bypass |
| D32 | CI, AS, BRB | | FC | | | 1.57 | 114 | 36.8 | 38 | Female | Gastric bypass |
| D33 | CI, AS, BRB | | FC | | | 1.6 | 100 | 40.56 | 46 | Female | Nephrectomy |
| D34 | CI, AS, BRB | | FC | | | 1.75 | 56 | 21.87 | 48 | Female | Nephrectomy |
| D35 | CI, AS, BRB | | FC | | TW AS | 1.58 | 95 | 31.02 | 72 | Female | Nephrectomy |
| D36 | | | FC | | | | | | 50 | Female | Gastric bypass |
| D37 | | | FC | | | 1.59 | 114 | 45.09 | 27 | Female | Gastric bypass |
| D38 | | | FC | | | 1.85 | 169 | 49.37 | 40 | Male | Gastric bypass |
| D39 | | | FC | | | 1.56 | 102 | 41.91 | 44 | Female | Gastric bypass |
| D40 | | | FC | | | 1.83 | 176.8 | 52.79 | 30 | Male | Gastric bypass |
| D41 | | | FC | CI, AS | | 1.66 | 118 | 42.82 | 42 | Female | Gastric bypass |
| D42 | | | FC | | | 1.73 | 106 | 35.41 | 55 | Male | Gastric bypass |
| D43 | | | FC, AS | | | 1.65 | 98 | 35.99 | 21 | Female | Gastric bypass |
| D44 | | | FC, AS | CI, AS | | | | | 48 | Male | Gastric bypass |
| D45 | | | FC | | | 1.79 | 110 | 34.33 | 40 | Male | Gastric bypass |
| D46 | | | FC | | | 1.67 | 90 | 32.27 | 44 | Male | Gastric bypass |
| D47 | | | FC | | | 1.64 | 122 | 45.35 | 45 | Male | Gastric bypass |
| D48 | | | FC | | | 1.71 | 133 | 45.48 | 34 | Female | Gastric bypass |
| D49 | | | FC | | | 1.47 | 85 | 39.33 | 53 | Female | Gastric bypass |
| D50 | | | FC | | | 1.67 | 105 | 37.64 | 56 | Female | Gastric bypass |
| D51 | | | FC | | | 1.85 | 175 | 51.13 | 34 | Male | Gastric bypass |
| D52 | | | FC | | | 1.7 | 141 | 48.78 | 22 | Female | Gastric bypass |
| D53 | | | FC | | TW AS, CI | 1.71 | 136 | 46.51 | 53 | Male | Gastric bypass |
| D54 | | | FC | | TW AS | 1.64 | 86 | 31.97 | 43 | Female | Gastric bypass |
| D55 | | | FC | | | 1.64 | 136 | 50.56 | 53 | Female | Gastric bypass |
| D56 | | | FC | | | 1.64 | 129 | 47.96 | 30 | Female | Gastric bypass |
| D57 | | | FC | | | 1.7 | 127 | 43.94 | 54 | Male | Gastric bypass |
| D61 | | scRNAseq | FC | | | 1.76 | 81 | 26.14 | 75 | Male | Nephrectomy |
| D62 | | | FC | | Chem AS, CI | 1.72 | 85 | 28.73 | 59 | Female | Nephrectomy |
| D63 | | | FC | | Chem AS, CI | 1.61 | 84 | 32.4 | 48 | Female | Nephrectomy |
| D64 | | | FC | | | 1.65 | 78 | 28.65 | 33 | Female | Nephrectomy |
| D65 | | | FC, AS | | TW AS, CI, ELISA | 1.73 | 88 | 29.4 | 66 | Male | Nephrectomy |
| D66 | | | FC, AS | | TW AS, ELISA | 1.65 | 65 | 23.87 | 44 | Female | Nephrectomy |
| D67 | | | FC, AS | Mix AS, IHC, BF | | 1.59 | 73 | 28.87 | 67 | Female | Nephrectomy |
| D68 | | | FC, AS, CI | Mix CI, AS | | 1.82 | 78 | 23.54 | 27 | Male | Nephrectomy |
| D69 | | | FC, AS | | | 1.62 | 70 | 26.67 | 59 | Female | Nephrectomy |
| D70 | | | FC, AS, CI | | | 1.76 | 83 | 26.79 | 59 | Male | Nephrectomy |
| D71 | | | FC | | | 1.77 | 83 | 26.49 | 42 | Male | Nephrectomy |
| D72 | | | | AS | | 1.77 | 129 | 41.17 | 70 | Male | Gastric bypass |
| D73 | | | | | Chem AS, CI | 1.7 | 80 | 27.68 | 61 | Female | Nephrectomy |
| D74 | | | | | KD, Chem AS | 1.5 | 71 | 31.55 | 55 | Female | Nephrectomy |
| D75 | | | | | Chem AS, KD, ELISA | 1.86 | 76 | 21.99 | 55 | Male | Nephrectomy |
| D76 | | | | | Chem AS, ELISA | 1.72 | 91 | 30.75 | 38 | Male | Nephrectomy |

**Legend**

| | |
|---|---|
| AS | Adiposcore |
| BF | brightfield images |
| BRB | Barcoded bulkRNA sequencing |
| Chem | chemical experiments |
| CI | Confocal imaging |
| ELISA | enzyme-linked immunosorbent assay |
| FC | flow cytometry |
| IHC | Immunohistochemistry |
| KD | siRNA knockdown experiments |
| Mix | mixing experiments |
| Profiles | flow cytometry scatter plot profiles |
| scRNAseq | single-cell RNA sequencing |
| TW | transwell |

### Supplementary Table 1 – Donor's information

| | Cohort of obese patients | Control cohort of kidney donors |
|---|---|---|
| Number | 55 | 26 |
| Gender (M:F) | 21:34 | 8:17 |
| Age Mean ± SD | **42.4** ± 11.7 | **53.9** ± 13.6 |
| BMI Mean ± SD [kg/m2] | **43.4** ± 6.6 | **26.8** ± 3.3 |
| Weight Mean ± SD [kg] | **124.1** ± 25.1 | **75.3** ± 12.2 |
| Height Mean ± SD [cm] | **168** ± 9 | **167.3** ± 9.3 |
| Type of surgery | Gastric bypass | Live donor nephrectomy |
| Accessible biopsies | SC, OM, MC | SC, OM, PR |

**Supplementary Table 2 – Cohorts specifications**

| Donor code | Age | Gender | Weight [kg] | Height [m] | BMI [kg/m2] | Type of surgery | Enrichment for SVF Lin– before scRNAseq | 10x Chromim 3' kit version | Analysed depots |
|---|---|---|---|---|---|---|---|---|---|
| D00 | 32 | Female | 138 | 1.74 | 45.58 | Gastric bypass | FACS | V2.0 | SC, OM |
| D01 | 46 | Female | 110 | 1.77 | 35.11 | Gastric bypass | FACS | V2.0 | SC, OM |
| D07 | 51 | Male | 128 | 1.79 | 39.94 | Gastric bypass | MACS | V3.1 | SC, OM, MC1 and MC2 |
| D24 | 63 | Female | 57 | 1.53 | 24.34 | Live donor nephrectomy | FACS | V3.1 | PR |
| D30 | 63 | Female | 60 | 1.68 | 21.25 | Live donor nephrectomy | FACS | V3.1 | PR |
| D61 | 75 | Male | 81 | 1.76 | 26.14 | Live donor nephrectomy | FACS | V3.1 | PR |

**Supplementary Table 3 – Donor’s specifications for scRNA-seq**

Red: Obese, Green: Normoweight, Yellow: Overweight

| Target | Provider | Catalog number | Host specie | Fluorophore conjugate | Biotium Mix n Stain fluorofore conjugate | Biotium Mix n Stain reference | Working titration | SC SVF Panel | OM SVF Panel | PR SVF Panel |
|---|---|---|---|---|---|---|---|---|---|---|
| Human HHIP | Sigma Aldrich | WH0064399M1 | Mouse | uncoupled | CF568 | 92235 | 1:20 | | | |
| Human MSLN | Biolegend | 530101 | Mouse | uncoupled | CF568 | 92255 | 1:25 | | | |
| Human VAP-1 | R&D | IC39571G | Mouse | AF488 | - | - | 1:80 | | | |
| Human CD26 | Biolegend | 302714 | Mouse | PE/Cy7 | - | - | 1:80 | | | |
| Human CD45 | Biolegend | 304022 | Mouse | Pacific Blue | - | - | 1:400 | | | |
| Human CD31 | Biolgend | 303114 | Mouse | Pacific Blue | - | - | 1:100 | | | |
| Human TM4SF1/L6 | R&D | FAB8164R | Mouse | AF647 | - | - | 1:20 | | | |

**Supplementary Table 4 – Antibody specifications for FACS**

| Target | Provider | Catalog number | Host specie | Fluorophore conjugate | Biotium Mix n Stain fluorofore conjugate | Biotium Mix n Stain reference | Working titration | Primary (I) or Secondary (II) |
|---|---|---|---|---|---|---|---|---|
| Human MSLN | Biolegend | 530101 | Mouse | uncoupled | CF568 | 92255 | 1:50 | I |
| Human TM4SF1/L6 | R&D | FAB8164R | Mouse | AF488 | - | - | 1:50 | I |
| Human PLIN1 | Abcam | ab172907 | Rabbit | uncoupled | | | 1:200 | I |
| anti-Rabbit IgG (H+L) | Thermo | A-31573 | Donkey | AF647 | | | 1:200 | II |

**Supplementary Table 5 – Antibody specifications for IHC**
Grey indicates secondary antibody
